# Neuroligin-3 interaction with CSPG4 regulates normal and malignant glial precursors through PIEZO1

**DOI:** 10.1101/2025.07.12.664340

**Authors:** Shawn M. Gillespie, Yoon Seok Kim, Anna C. Geraghty, Belgin Yalçın, Rebecca Mancusi, Jared Hysinger, Alexis English Ivec, James Reed, Richard Drexler, Michael Quezada, Karen Malacon, Pamelyn Woo, Christopher Mount, Aerin Yang, Mable Lam, Yuan Pan, J. Bradley Zuchero, Jacqueline Trotter, Michelle Monje

**Affiliations:** Department of Neurology, Stanford University, Stanford, CA, USA; Department of Molecular and Cellular Physiology, Stanford University, Stanford, CA, USA; Department of Neurosurgery, Stanford University, Stanford, CA, USA; Department of Symptom Research, MD Anderson Cancer Center, Houston, TX, USA; Department of Biology, Molecular Cell Biology, Johannes Gutenberg University, Mainz, Germany; Department of Psychiatry and Behavioral Sciences, Stanford University, Stanford, CA, USA; Department of Pathology, Stanford University, Stanford, CA, USA; Department of Pediatrics, Stanford University, Stanford, CA, USA; Institute for Stem Cell Biology and Regenerative Medicine, Stanford University, Stanford, CA, USA; Howard Hughes Medical Institute, Stanford, CA, USA

## Abstract

Glioma pathophysiology is robustly regulated by interactions with neurons. Key to these interactions is the role of neuroligin-3 (NLGN3), a synaptic adhesion molecule shed in response to neuronal activity^1–5^ that functions as a paracrine factor crucial for glioma growth. Here, we elucidate the mechanistic pathway whereby shed NLGN3 interacts with glioma and their normal glial counterpart. NLGN3 interacts with Chondroitin Sulfate Proteoglycan 4 (CSPG4) on both glioma and healthy oligodendrocyte precursor cells (OPCs)^6–9^, facilitating CSPG4 shedding by ADAM10. NLGN3-CSPG4 interactions and consequent shedding alter membrane tension, thereby activating PIEZO1 mechanosensitive channels and causing membrane depolarization. The NLGN3-CSPG4-PIEZO1 axis maintains OPCs in an undifferentiated, stem-like state and promotes glioma proliferation, underscoring important functional roles for the NLGN3-CSPG4-PIEZO1 axis in both healthy and malignant glial precursors.

---

Gliomas are a devastating family of brain cancers. High-grade gliomas such as glioblastoma (GBM) and H3K27M-altered diffuse midline gliomas (DMG, also called diffuse intrinsic pontine glioma (DIPG)) are the leading cause of primary brain tumor-related death in both adults and children, while low-grade gliomas cause substantial neurological morbidity. Many forms of glioma arise from oligodendrocyte precursor cells (OPC), and OPC-like cancer cells comprise an important cellular subpopulation within gliomas such as glioblastoma and diffuse midline glioma (for review, please see Taylor and Monje, 2023)^10^. Concordantly, many mechanistic parallels exist between healthy OPCs and malignant gliomas. A key similarity is the proliferation-promoting effect of neuronal activity. Glutamatergic neuronal activity robustly promotes the proliferation of both healthy OPCs^11^ and glioma cells^1^. In OPCs, this activity-regulated proliferation results in the generation of new mature oligodendrocytes and adaptive changes in myelination that tune neural circuit function and contribute to brain functions like attention, learning and memory^11–14^. Gliomas hijack these mechanisms of glial plasticity to promote growth^1–3,15,16^ and invasion^17,18^.

Neuron-glioma interactions are mediated by activity-regulated paracrine growth factors^1–3^ and by electrophysiologically functional neuron-to-glioma synapses^15,16^. One important mechanism underlying activity-regulated glioma growth is the activity-regulated shedding of the synaptic adhesion molecule neuroligin-3 (NLGN3)^1,2^. Genetic mouse modeling experiments conditionally knocking out NLGN3 in a cell type-specific manner demonstrated that OPCs are the major cellular source of shed NLGN3^2^. For context, it is important to note that OPCs receive extensive synaptic inputs from neurons^19–21^, although the function of neuron-to-OPC synapses is still incompletely understood. NLGN3 shedding occurs through neuronal activity regulation of ADAM10 protease function, which cleaves NLGN3 at the membrane to release the large N-terminal ectodomain into the extracellular microenvironment. Shed NLGN3 promotes the proliferation of both high-grade^1,2,22^ and low-grade^3^ gliomas through stimulation of proliferation signaling pathways PI3K-mTOR^1,2^, RAS and SRC^2^. Unexpectedly, genetic knockout of *Nlgn3* gene from the cells in the microenvironment or pharmacological blockade via ADAM10 inhibition of Nlgn3 shedding in the brain microenvironment markedly slows glioma growth and progression in preclinical models of high-grade gliomas (adult glioblastoma, pediatric glioblastoma, diffuse intrinsic pontine glioma (DIPG)/DMG)^2^ and low-grade glioma (NF1-associated optic pathway glioma)^3^. The stark dependency of gliomas on NLGN3 signaling in these experimental models can be understood in part by observations that NLGN3 exposure increases glioma gene expression related to multiple processes of neuron-glioma interactions, notably synapse-related gene expression^2^ and concordantly neuron-to-glioma synapses are reduced in the absence of NLGN3 from the tumor microenvironment^16^. Taken together, these observations highlight the therapeutic potential of disrupting NLGN3 shedding or downstream effects on glioma cells. However, a number of pressing questions remain that include mechanisms by which NLGN3 interacts with and signals to the glioma cell, whether similar interactions occur on healthy OPCs, and what role NLGN3 shedding may play in healthy oligodendroglial biology. Here, we aim to address these questions to elucidate the mechanism of NLGN3 signaling to OPCs and glioma cells.

## Unbiased Biochemical Screening for NLGN3 binding partners

To screen for putative interaction partners of shed NLGN3, we leveraged affinity capture chromatography using a NLGN3 column^23,24^. We first produced recombinant NLGN3 ectodomains (amino acids 1-689) and biotinylated them via a C-terminal Avitag. We then immobilized the NLGN3 ectodomains on commercial streptavidin resin (Fig. 1a and b) Next, we isolated and solubilized membranes from patient-derived cell cultures of DIPG/DMG or adult GBM (Fig. 1a) and passed each glioma membrane sample through the column in separate experiments. After washing, the eluate was separated by SDS/PAGE and analyzed by mass spectrometry (Materials and Methods.). We identified several putative NLGN3 interacting partners from both cell cultures, prominently including Chondroitin Sulfate Proteoglycan 4 (CSPG4) (Fig. 1b). Of note, the N-terminus of CSPG4 (also known as NG2 in rat, AN2 in mouse) contains two laminin-neurexin-sex-hormone binding globulin (LNS) domains (Fig. 1c)^25–27^. Similar LNS domains in pre-synaptic neurexins participate in their well-characterized interaction with post-synaptic neuroligins^28,29^, highlighting CSPG4 as a likely interaction partner. Of note, CSPG4 (NG2) is a canonical marker of OPCs (which were previously referred to as “NG2 cells”) and is also highly expressed in a variety of gliomas, particularly in the OPC-like cellular compartment of glial malignancies.

**Fig. 1.**
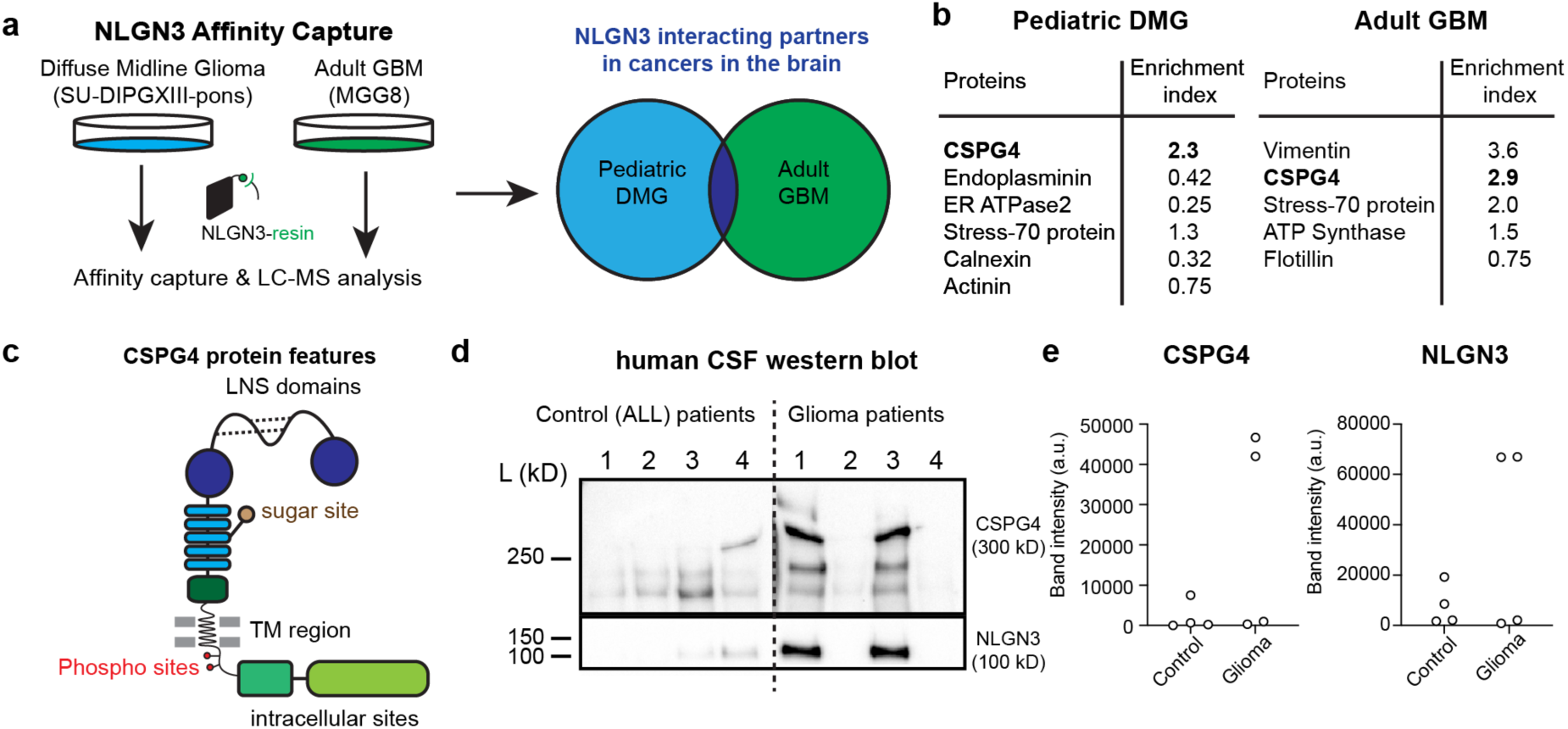
Unbiased screening identifies CSPG4 as a putative interacting partner of shed NLGN3. **a.** Schematic of glioma membrane isolation, solubilization, and affinity capture by immobilized NLGN3 ectodomains. NLGN3 affinity column is prepared by immobilizing biotinylated (c- terminal AviTag) NLGN3 ectodomains within a streptavidin resin **b.** Selected top hits from the NLGN3 affinity chromatography assay. Purified membrane proteins captured through NLGN3 affinity chromatography were analyzed by mass spectrometry. Note that both pediatric (DIPGXIII) and adult (MGG8) brain cancers show CSPG4 as a top hit. **c.** Cartoon representation of CSPG4 functional domains. The laminin-G (LNS) domains of CSPG4 are analogous to those of Neurexins (NRXNs), the canonical binding partners to NLGN3, followed by the signature chondroitin sugar-binding site in the middle of core repeat modules. **d.** Western blot of human CSF with anti-CSPG4 N-term and anti-NLGN3 antibodies. Control patient CSFs are all from patients with acute lymphocytic leukemia (ALL). Only CSF samples from diffuse midline glioma-bearing patients show both CSPG4 and NLGN3. **e.** Quantification of bands in d (n = the number of patients = 4/group, mean ± s.e.m).

To quantitatively verify that NLGN3 shows dose-dependent binding to chondroitin sulfate (CS), the sugar substrate that is covalently linked to CSPG4 (Fig. 1c), we performed a well-established fluorescent size-exclusion chromatography (FSEC) assay (Extended Data Fig. 1a) using recombinant NLGN3 ectodomain with CS and the established NLGN-interacting partner, the glycosaminoglycan heparin^30,31^. Both glycans showed a significant left shift in the FSEC profile, indicating that the molecular weight of NLGN3 increased due to the binding of the sugar substrate (Extended Data Fig. 1b). Although the shift caused by CS binding was less than that of heparin, it showed robust dose-dependent binding (Extended Data Fig. 1c, d), suggesting that the CS is likely a direct binding substrate of NLGN3. Taken together, our biochemistry data strongly support our hypothesis that CSPG4 is an interacting partner for NLGN3.

CSPG4 ectodomains are also reported to be shed in response to neuronal activity^9^, similar to NLGN3. Therefore, we sought to determine whether shed ectodomains of NLGN3 and CSPG4 are detectable in cerebrospinal fluid (CSF) of human glioma patients. We obtained CSF from four patients with DIPG/DMG and four patients with leukemia (negative control group) and performed western blot analysis with antibodies against the N-termini of NLGN3 and CSPG4 (Fig. 1d). In alignment with prior findings of NLGN3 shedding from glioma cells^2^, we readily detected both NLGN3 and CSPG4 protein in CSF from two of four glioma patients, and none of the leukemia patients (Fig. 1e). Altogether, these data support CSPG4 as a candidate interacting partner for shed NLGN3, but do not clarify how these proteins interact, or if their interaction is related to their shedding in the context of neuronal activity.^1–3,32^.

## NLGN3 exposure augments ADAM10-mediated CSPG4 proteolysis

Previous work demonstrated that neuronal activity triggers ADAM10-mediated cleavage of NLGN3 ectodomains from OPC, glioma, and, to a lesser extent, neuronal cell membranes^2^. Similarly, CSPG4 undergoes ADAM10-mediated ectodomain cleavage in response to neuronal activity^9^. Here, we find that while NLGN3 and CSPG4 ectodomains are both shed from membranes in response to neuronal activity, these events are separated in space and time. We hypothesized that shed NLGN3 subsequently triggers CSPG4 shedding from the membrane of glioma cells and OPCs in an autocrine and/or paracrine fashion. To directly assess this hypothesis, we treated *in vitro* cell cultures of glioma cells with recombinant NLGN3 ectodomain proteins for one hour, collected the conditioned medium, and performed western blot analysis using antibodies against the CSPG4 N-terminus (Fig. 2a).

**Fig. 2.**
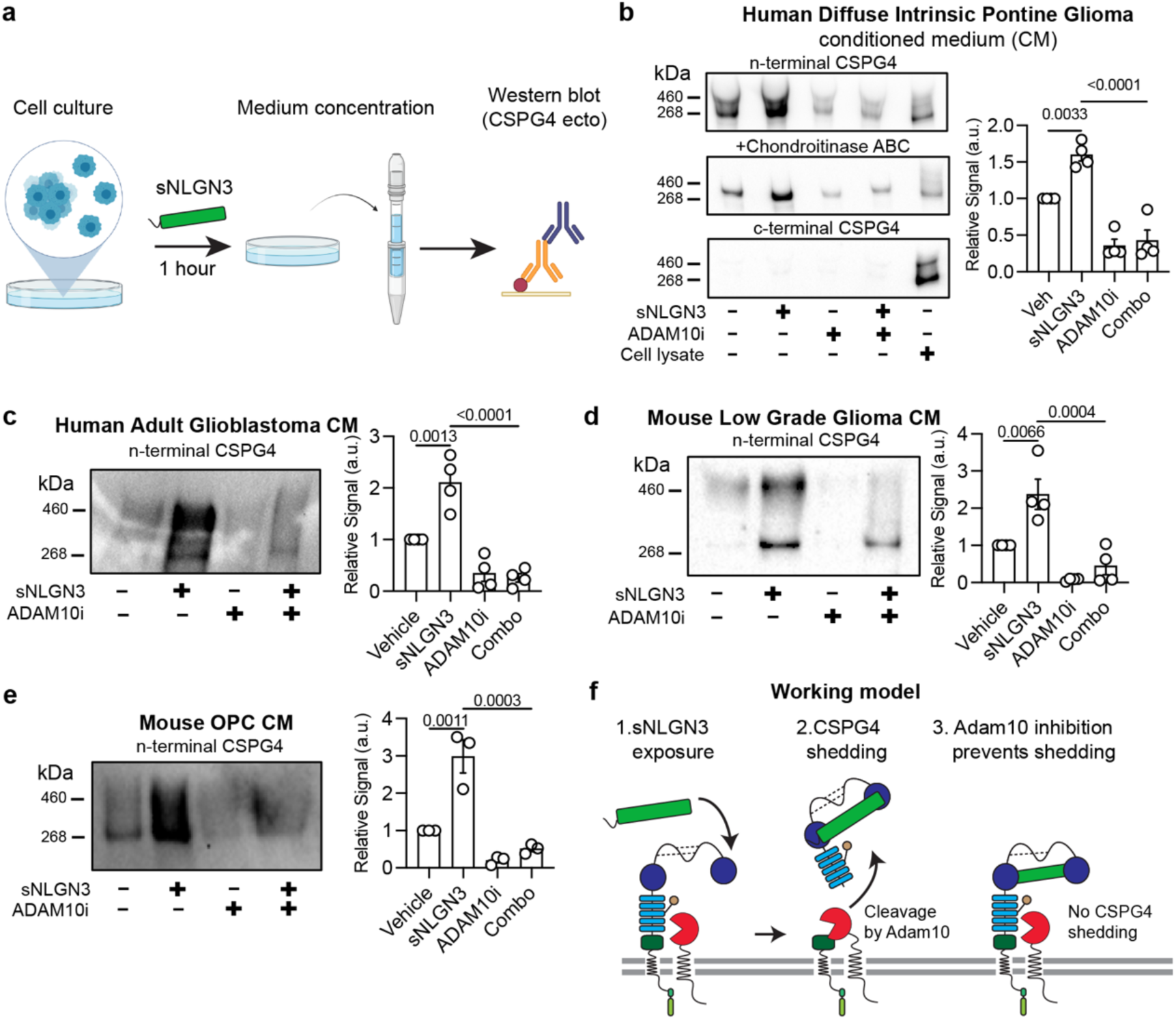
Shed NLGN3 treatment augments ADAM10-dependent CSPG4 cleavage. **a.** Workflow depicting treatment of glioma cells or OPCs for 1 hour with recombinant NLGN3 ectodomains (to mimic naturally shed NLGN3) followed by conditioned medium (CM) concentration and western blot to detect shedding of CSPG4. **b.** *<u>Left</u>*: CSPG4 Western blot from diffuse intrinsic pontine glioma model SU-DIPG13 conditioned medium after exposure to 100 nM sNLGN3 for 1-hour in the presence or absence of 5 uM ADAM10 inhibitor. CSPG4 exists as a ‘part-time’ proteoglycan and is detected as either a core protein (discrete band at ∼268 kDa) or as a chondroitin sulfate proteoglycan (higher molecular weight smear). Treatment of conditioned medium with chondroitinase ABC removes chondroitin sulfate chains and collapses diffuse smears into discrete bands. A c-terminal specific CSPG4 antibody only detects protein in the cell lysate control lane, confirming that conditioned medium bands are shed ectodomains. NLGN3 treatment increases shedding relative to vehicle controls (p = 0.003). NLGN3-induced shedding of CSPG4 is blocked by pre-treating cells with 5 uM ADAM10 inhibitor (p = <0.0001). *Right*: Quantification of CSPG4 shedding as fold change in protein band intensities in the left (top N-terminal blot) relative to vehicle-treated controls. n (independent biological replicates of western blot) = 4, mean ± s.e.m., ordinary one-way ANOVA, p-values are mentioned in the legend and written on the figures. **c.** *<u>Left</u>*: Western blot of CSPG4 shedding from IDH1 WT glioblastoma (GBM) model conditioned medium after exposure to 100 nM sNLGN3 for 1-hour in the presence or absence of 5 uM ADAM10 inhibitor. NLGN3 treatment increases shedding relative to vehicle controls (p = 0.0013). NLGN3-induced shedding of CSPG4 is blocked by pre-treating cells with 5 uM ADAM10 inhibitor (p = <0.0001). *<u>Right</u>*: Quantification of protein band intensities relative to vehicle-treated controls. n (independent biological replicates of western blot) = 4, mean ± s.e.m., ordinary one-way ANOVA, p-values are mentioned in the legend and written on the figures. **d.** *<u>Left</u>*: Western blot of CSPG4 shedding from NF1-mutant optic pathway glioma model conditioned medium after exposure to 100 nM sNLGN3 for 1-hour in the presence or absence of 5 uM ADAM10 inhibitor. NLGN3 treatment increases shedding relative to vehicle controls (p = 0.0066). NLGN3-induced shedding of CSPG4 is blocked by pre-treating cells with 5 uM ADAM10 inhibitor (p = 0.0004). *<u>Right</u>*: Quantification of protein band intensities relative to vehicle-treated controls. n (independent biological replicates of western blot) = 4, mean ± s.e.m., ordinary one-way ANOVA, p-values are mentioned in the text and the figures. **e.** *<u>Left</u>*: Western blot of CSPG4 shedding from primary mouse oligodendrocyte precursor cell (OPC) conditioned medium after exposure to 100 nM sNLGN3 for 1-hour in the presence or absence of 5 uM ADAM10 inhibitor. NLGN3 treatment increases shedding relative to vehicle controls (p = 0.0011). NLGN3-induced shedding of CSPG4 is blocked by pre-treating cells with 5 uM ADAM10 inhibitor (p = 0.0003) *<u>Right</u>*: Quantification of protein band intensities relative to vehicle-treated controls. n (independent biological replicates of western blot) = 3, mean ± s.e.m., ordinary one-way ANOVA, p-values are mentioned in the figures. **f.** Model depicting serial events occurring during NLGN3-induced CSPG4 ectodomain shedding by ADAM10 from glioma cells. Red circular sector = ADAM10; Green rectangle = shed NLGN3; blue = CSPG4 ectodomain.

We observed that glioma cells constitutively shed CSPG4 into their culture medium (Fig. 2b), a process inhibited by brief treatment with ADAM10 inhibitor GI254023X (5 µM, 15 mins) (Fig. 2b). Notably, NLGN3 treatment led to a significant increase in CSPG4 cleavage (Fig. 2b), which was also inhibited by treatment of cells with ADAM10 inhibitor (Fig. 2b). Importantly, we did not detect the C-terminus of CSPG4 from conditioned medium, confirming the absence of cytoplasmic domains in these shed ectodomains (Fig. 2b). Additionally, we found that DIPG/DMG-derived CSPG4 ectodomains exhibited variable glycosylation, as treatment of the conditioned medium with CS-lysing enzyme chondroitinase ABC collapsed diffuse smears into discrete bands in the gel (Fig. 2b).

Expanding beyond high-grade pediatric gliomas such as DIPG/DMG, we established that NLGN3-induced CSPG4 shedding occurs across a spectrum of glioma subtypes, including human IDH1 WT adult glioblastoma (Fig. 2c) and a mouse model of NF1-associated optic pathway glioma^3^ (Fig. 2d). Furthermore, we confirmed that this mechanism is not exclusive to malignant cells, as primary mouse OPCs readily shed CSPG4 ectodomains in response to recombinant NLGN3 (Fig. 2e). Consistent with previous findings^9^, NLGN3-induced shedding of CSPG4 was inhibited by pre-treating OPCs with an ADAM10 inhibitor. These results collectively demonstrate that NLGN3 exposure is sufficient to enhance ADAM10-dependent shedding of CSPG4 ectodomains from OPCs and glioma cells (Fig. 2f).

## Membrane depolarization disinhibits ADAM10 and increases CSPG4 shedding

We next aimed to clarify the mechanism by which NLGN3 exposure augments ADAM10- dependent CSPG4 ectodomain shedding. It is known that ADAM10-mediated shedding of transmembrane protein substrates, like NLGN3 and CSPG4, is regulated in part by an auto- inhibitory c-loop mechanism that obstructs the active site^33^. This inhibitory effect is relieved upon cellular depolarization, which leads to an imbalance of charge between the inner and outer leaflets of the plasma membrane, consequently exposing the active site and permitting substrate cleavage^34^. To investigate whether this mechanism occurs in glioma cell membranes, we conducted a CSPG4 shedding assay in which cells were exposed to various depolarizing stimuli in comparison to NLGN3 treatment (Fig. 3)^9^.

**Fig. 3.**
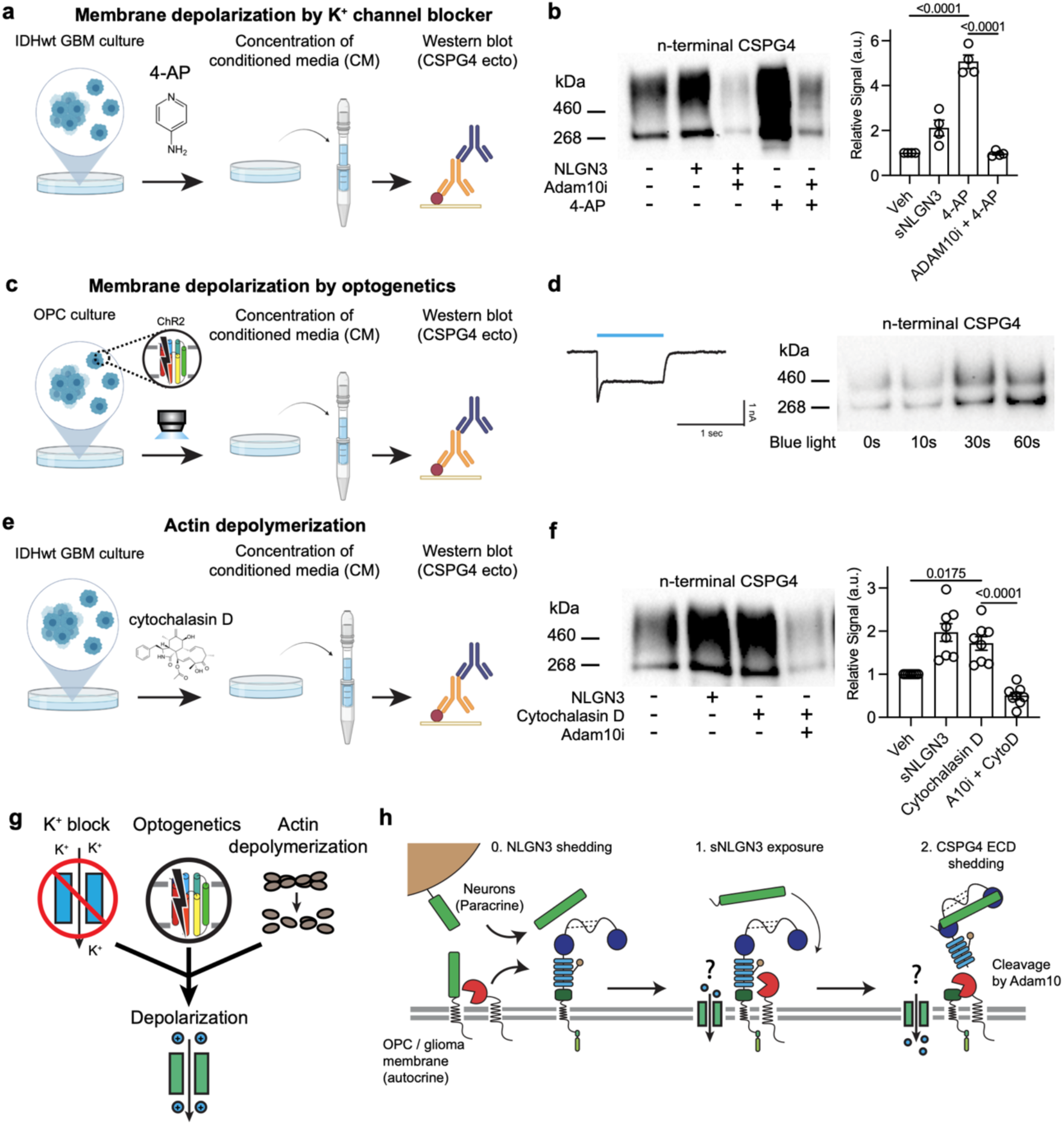
Membrane depolarization and actin depolymerization regulate ADAM10- dependent CSPG4 shedding. **a.** Workflow depicting treatment of IDH1 WT GBM (MGG8) cells for 1 hour with the potassium channel blocker 4-Aminopyridine (4-AP) followed by culture medium concentration and western blot to detect shedding of CSPG4. **b.** *<u>Left</u>*: Representative western blot of shed CSPG4 ectodomains from experiment depicted in a. 4-AP treatment increases shedding relative to vehicle controls. Note that 4-AP-induced shedding of CSPG4 is blocked by pre-treating cells with 5 uM ADAM10 inhibitor (n (independent biological replicates of western blot) = 4, mean ± s.e.m., ordinary one-way ANOVA, p < 0.0001) *<u>Right</u>*: Quantification of western blots. (n (independent biological replicates of western blot) = 4, mean ± s.e.m., ordinary one-way ANOVA, p-values are written on the figures. **c.** Workflow depicting blue light stimulation of Channelrhodopsin-2 (ChR2)-expressing rat OPCs followed by culture medium concentration and western blot to detect shedding of CSPG4. **d.** *Left:* Representative current of ChR2-expressing OPCs. (Light power density = 1.0 mW/mm^2^, wavelength = 470 nm, duration = 1 second). *Right:* Representative western blot of shed CSPG4 ectodomains from experiment depicted in c (pulse width = 25 ms, frequency = 10 Hz). Membrane depolarization by ChR2 stimulation increases shedding relative to no blue light stimulation controls. **e.** Workflow depicting treatment of IDH1 WT GBM cells (MGG8) for 1-hour with the actin polymerization inhibitor cytochalasin D followed by culture medium concentration and western blot to detect shedding of CSPG4. **f.** *<u>Left</u>*: Western blot of shed CSPG4 ectodomains from experiment depicted in e. Cytochalasin D treatment increases shedding relative to vehicle controls (p = 0.0175). Cytochalasin D-induced shedding of CSPG4 is blocked by pre-treating cells with 5 uM ADAM10 inhibitor (p < 0.0001) *<u>Right</u>*: Quantification of western blots (n (independent biological replicates of western blot) = 7, mean ± s.e.m., ordinary one-way ANOVA, p-values are written on the figures. **g.** Cartoon diagram summarizing the treatments of 4-AP, ChR2, and cytochalasin D, that are inducing membrane depolarization. **h.** Model depicting CSPG4 shedding through NLGN3 interaction that implicates an ion channel that is sensitive to membrane stress to depolarize the membrane and activate ADAM10.

To induce prolonged membrane depolarization, glioma cell cultures were treated with the voltage-gated K^+^ channel blocker, 4-aminopyridine (4-AP, 2.5 mM, 75 mins), a well-characterized reagent for enhancing signal transduction of electrically responsive cells in various experimental applications^35,36^. Following treatment with 4-AP, glioma cell conditioned medium was collected, concentrated and profiled by CSPG4 western blot (Fig. 3a). Akin to NLGN3 exposure, 4-AP strongly enhanced CSPG4 shedding. (Fig. 3b). Consistent with NLGN3 exposure, shedding induced by 4-AP was significantly diminished by co-treatment with ADAM10 inhibitor (Fig. 3b).

We next investigated whether ADAM10-mediated shedding of CSPG4 is contingent upon the duration of cellular membrane depolarization; to address this question, we employed optogenetics—a method developed for precise spatiotemporal control of cellular membrane potential using light stimuli^37^. We achieved this by expressing the light-gated cation channel channelrhodopsin-2 (ChR2) in OPCs through lentiviral transduction and subjecting them to blue light stimulation (Fig. 3c) (470 nm, 1 mW/mm^2^, 0-60 seconds) to induce transient and controlled depolarization (Fig. 3c, left). We observed a robust increase in CSPG4 shedding with optogenetic depolarization of OPCs (Fig. 3c, right). Notably, the magnitude of shedding was contingent upon the duration of ChR2 stimulation (Fig. 3c), indicating that depolarization directly enhanced ADAM10-mediated shedding of CSPG4. This finding strongly supports the regulation of ADAM10-mediated CSPG4 shedding by membrane depolarization.

Intriguingly, CSPG4 ectodomains are also reportedly shed in response to an actin depolymerizing agent^25,32,38^. The disruption of actin microfilaments can induce significant membrane depolarization^39,40^. Building on this prior literature, we reasoned that the CSPG4 shedding induced by actin depolymerizing could result from a membrane depolarization and consequent ADAM10 disinhibition. To test this hypothesis, we treated glioma cells with cytochalasin D, an actin depolymerase (2 µM, 15 minutes) before adding NLGN3 and performing a western blot to detect CSPG4 ectodomains in conditioned medium (Fig. 3e). We confirmed that the disruption of actin filaments indeed resulted in a notable increase in CSPG4 shedding (Fig. 3f). Importantly, cytochalasin D-induced shedding was also blocked by an ADAM10 inhibitor. Collectively, our results from these various strategies demonstrate that membrane depolarization enhances ADAM10-mediated shedding of NLGN3 (Fig. 3g). These findings also suggest that NLGN3 exposure results in a membrane depolarization that enhances ADAM10-mediated CSPG4 shedding (Fig. 3h). We therefore wondered if the membrane-bound actuators responsible for translating NLGN3 exposure into enhanced ADAM10 activity in glioma cells and OPCs were cation-conducting channels sensitive to disruptions in the actin cytoskeleton.

## NLGN3 disinhibits ADAM10 via PIEZO1 activation

Actin depolymerization, which mimics NLGN3 treatment in terms of ADAM10 activation and CSPG4 cleavage, is known to increase shear stress and membrane tension^41–43^. To explore whether NLGN3 exposure itself alters membrane tension, we first assessed changes in membrane tension using a membrane-infiltrating chemical dye Flipper-TR (Spirochrome), a widely used method to measure changes in membrane tension in live cells (Fig. 4a)^44,45^. Remarkably, we observed a significant increase in membrane tension in glioma cells after treatment with NLGN3 (Fig. 4a, left, Fig. 4b, left). Importantly, this increase was dependent on the presence of CSPG4 as we did not observe changes in membrane tension in CSPG4 KO glioma cells. (Fig. 4a, right, Fig. 4b, right). Given these results, we formulated a model in which the force of NLGN3 interacting with CSPG4 causes increased membrane tension that is sensed by mechanosensitive ion channels, depolarization-mediated disinhibition of ADAM10, and ultimately shedding of CSPG4 ^34,41–43,46,47^.

**Fig. 4.**
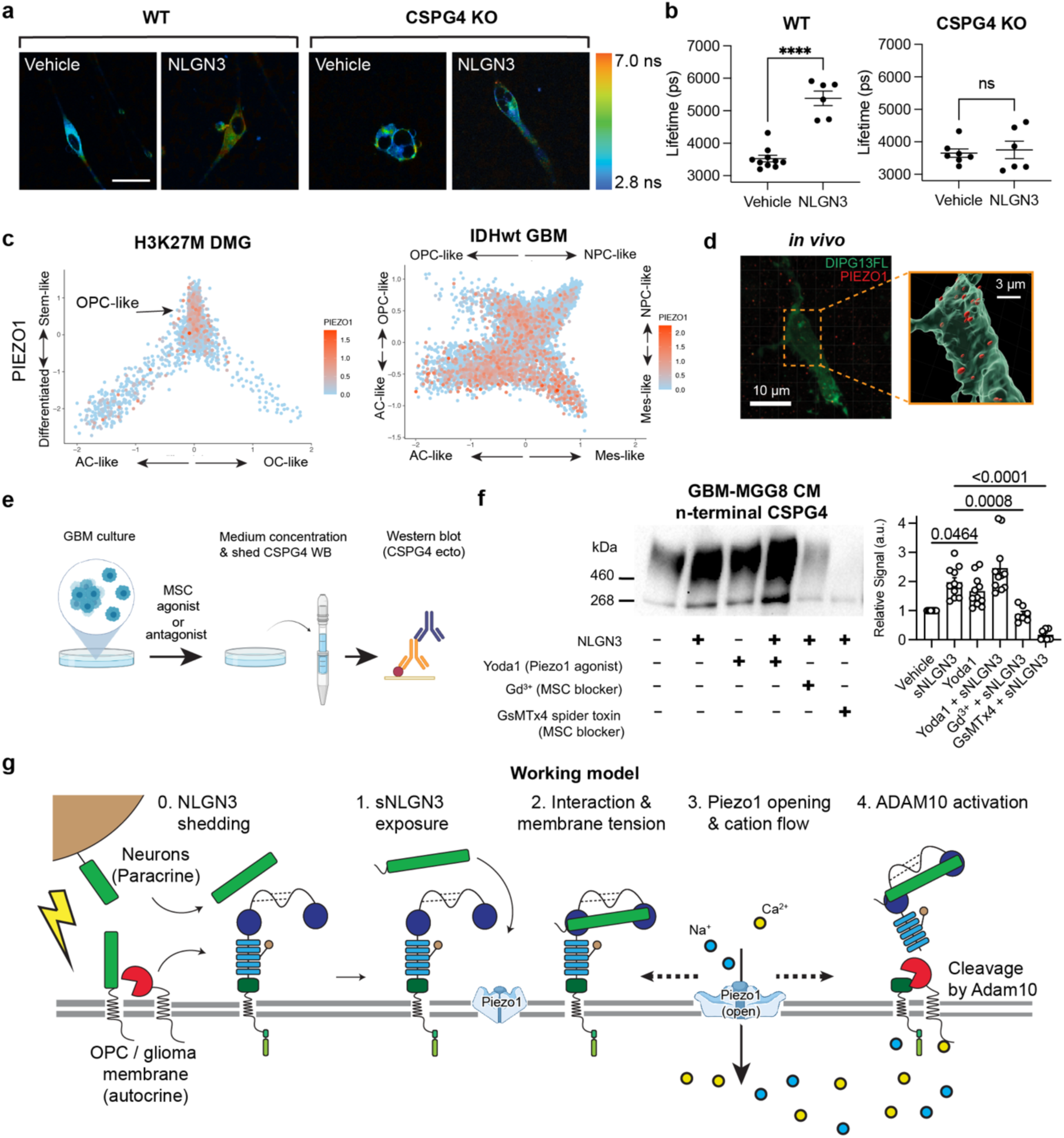
sNLGN3 induces ADAM10-dependent CSPG4 cleavage via activation of PIEZO1 channels. **a.** Representative images of glioma cells colored based on the lifetime of fluorescent tension sensor FlippeR-TR. The fluorescence probe works by specifically targeting the plasma membrane, and the longer lifetime corresponds to increased membrane tension. **b.** Quantification of the lifetimes measured from the experiment shown in a. (n (number of independent biological samples used for imaging) = 6 – 10, mean ± s.e.m., two-tailed t-test, **** P < 0.0001, n.s. = not significant) **c.** Scatterplots depicting tumor architecture of scRNA-seq produced from molecularly distinct patient glioma samples. From left to right are Histone H3K27M mutated diffuse midline glioma (DMG) and IDH1 wild-type glioblastoma (GBM). PIEZO1 expression displayed in red across two classes of glioma. Note that PIEZO1 is detected within the OPC-like fraction of these distinct glioma types. **d.** left: Confocal image of immunostaining in SU-DIPGXIII-FL xenografted mouse brain slice. (PIEZO1 = red, GFP (marker for glioma cells) = green, scale bar = 10 µm) right: 3-dimensional reconstructions of glioma cell surface showing PIEZO1 localization. Patient-derived DIPG cells (SU-DIPGXIII-FL; nestin, blue) co-localize with PIEZO1 puncta (red). Scale bar, 3 µm. **e.** Workflow depicting treatment of glioma cells with recombinant NLGN3 ectodomains in the presence or absence of modulators of mechanosensitive ion channels (MSC) followed by culture medium concentration and western blot to detect shedding of CSPG4. **f.** Western blot of shed CSPG4 ectodomains from experiment depicted in c. Treatment with Yoda1 (PIEZO1 agonist), increases shedding relative to vehicle controls (p = 0.0464). In contrast, treating cells with MSC blockers gadolinium (Gd^3+^) or the spider toxin GsMTx4 prevent NLGN3-induced shedding of CSPG4 (p = 0.0008 and p < 0.0001, respectively) *<u>Right</u>*: Quantification of western blots (n (independent biological replicates of western blot) = 11, mean ± s.e.m., ordinary one-way ANOVA, p-values are written on the figures. **g.** Model depicting CSPG4 shedding via ADAM10 activation that is downstream of NLGN3 interaction and ensuing membrane depolarization through PIEZO1 channel activation. Red circular sector = ADAM10; Green rectangle = shed NLGN3; blue = CSPG4 ectodomain; light blue = PIEZO1.

To validate our model, we endeavored to identify actuators that could be responsible for sensing and responding to the interaction between shed NLGN3 and CSPG4 and the resulting increase in membrane tension. Increased shear stress and membrane tension are known to activate mechanosensitive ion channels such as PIEZO and transient receptor potential (TRP) channels that are associated with the cytoskeleton^41–43,46,47^. In light of these studies, we interrogated multiple single-cell RNAseq datasets from pediatric diffuse midline gliomas (DMG) and adult IDH1 wild-type glioblastomas (GBM) to curate a list of mechanosensitive ion channels that are expressed in OPC-like cells also expressing CSPG4 (Fig.4c, Extended Data Fig. 2a). Proteins meeting these criteria include PIEZO1, PIEZO2 and several members of the TRP channel family (Extended Data Fig. 2b)^48,49^.

Given that the expression level of the mechanosensitive ion channel PIEZO1 was the highest among all candidates (Extended Data Fig. 3b) we prioritized PIEZO1 for further investigation. Immunostaining (Fig. 4d) and western blot (Extended Data Fig.. 2c) analyses indicated high levels of PIEZO1 in glioma cells in both *in vivo* and *in vitro* conditions. To assess whether PIEZO1 mediates NLGN3-induced ADAM10 activation, we conducted a CSPG4 shedding assay on glioma cells pre-treated with either the non-specific mechanosensitive channel blocker gadolinium ion (Gd^3+^) or the PIEZO1-specific blocker spider toxin GsMTx4 (Fig. 4e, Extended Data Fig.. 2d)^50^. Remarkably, blockade of PIEZO1 conduction proved to be as effective as the ADAM10 inhibitor in preventing NLGN3-induced shedding of CSPG4 (Fig. 4f, Extended Data Fig.. 2e, f). In contrast, treatment of cells with the PIEZO1 agonist Yoda1 resulted in enhanced shedding of CSPG4 (Fig. 4f, Extended Data Fig. 2e, f), suggesting the necessity and sufficiency of PIEZO1 activation in the process of NLGN3-induced activation of ADAM10 cleavage and CSPG4 shedding (Fig.4g)^51^.

We next investigated PIEZO1 activity in glioma cells using electrophysiology techniques to record mechanosensitive ion currents. Employing a cell-attached patch clamp configuration, we measured the currents during a series of increasing negative pressures applied via the recording pipette (Fig. 5a)^50–52^. Initially, we observed that a sequence of mechanical stimulations resulted in small, but significant, inward currents (Fig. 5a, b). Importantly, these currents were markedly attenuated by the presence of the PIEZO1 blocker GsMTx4 (7.9 ± 1.9 pA vs 2.8 ± 0.23 pA, at - 40 mmHg) and augmented by the agonist Yoda1^52^ (7.9 ± 1.9 pA vs 23.6 ± 6.0 pA, at -40 mmHg), confirming that these currents are primarily elicited by the activation of the PIEZO1 channel (Fig. 5b, c). In further support of this hypothesis, we observed a significant increase of PIEZO1 currents in the presence of NLGN3 (Fig. 5b, c)(7.9 ± 1.9 pA vs 17.1 ± 5.0 pA, at -40 mmHg), similar to the level of increase induced by Yoda1. This finding is consistent with the increase in the membrane tension observed earlier (Fig. 4) and clearly indicates that PIEZO1 activity is directly modulated by the presence of NLGN3. To further validate this data, we conducted cell-attached patch clamp recordings in CSPG4 knockout glioma cells. Surprisingly, we observed minimal currents after a series of negative pressures with and without NLGN3, with no difference between the two groups (Extended Data Fig.3a and b,). This data suggests that PIEZO1 activity is influenced by the expression of CSPG4 in this cellular context, and may be modulated by the interaction between NLGN3 and CSPG4.

**Fig. 5.**
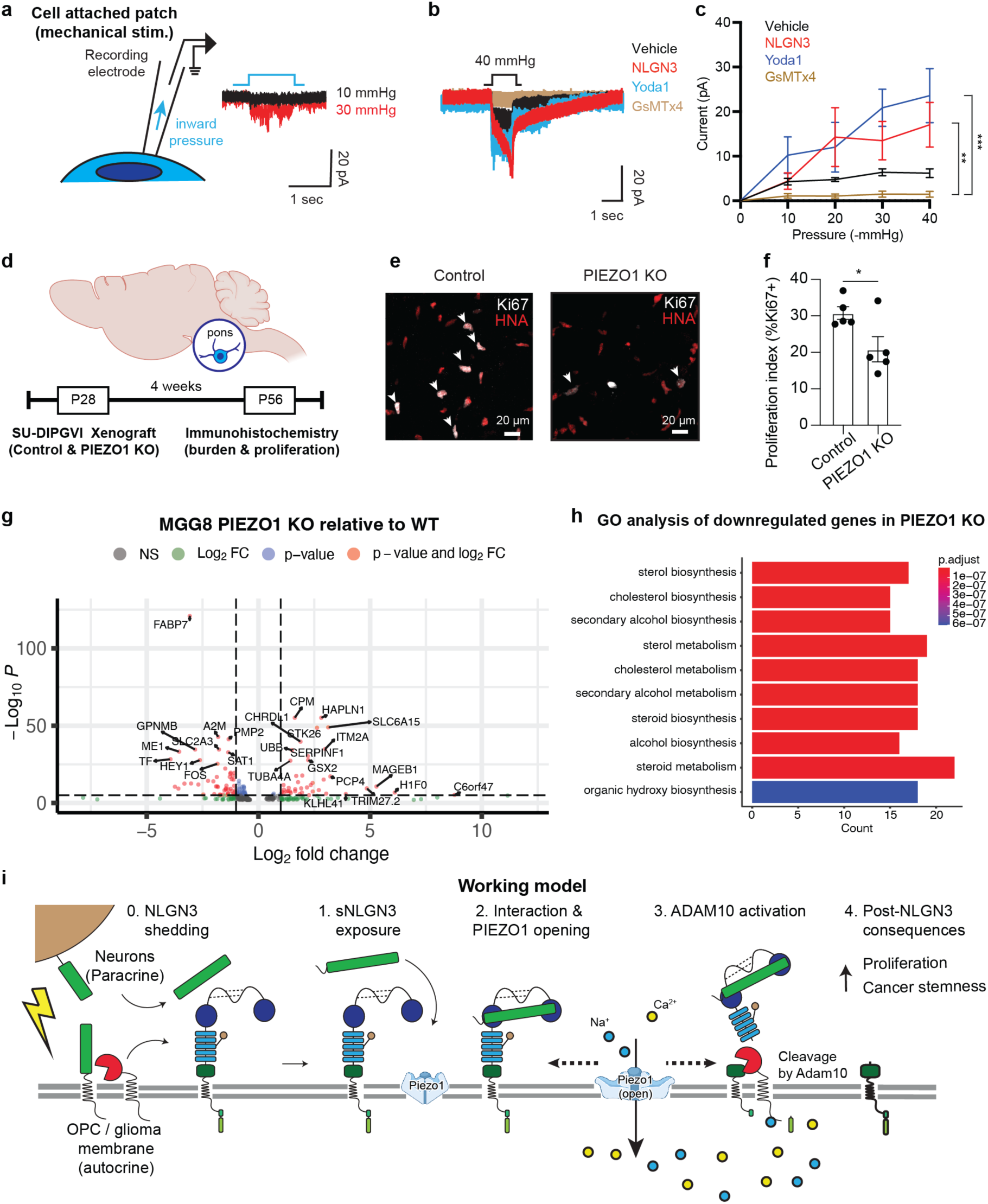
Blockade of CSPG4 shedding by PIEZO1 prevents NLGN3-induced changes to glioma cells. **a.** (Left) Schematics of cell-attached patch clamp recording of mechanosensitive currents from glioma cells. (Right) The representative waveforms of mechanosensitive currents measured with 10 mmHg and 30 mmHg pressure inputs. **b.** Representative traces of mechanosensitive currents (pressure input = 40 mmHg, 1 second) with vehicle (black), PIEZO1 agonist Yoda1 (cyan), NLGN3 (red) and PIEZO1 blocker GsMTx4 (brown). **c.** Summary of mechanosensitive currents shown in b. (n (number of independent biological samples) = 6 **–** 8, mean ± s.e.m., Kruskal-Wallis test, ** P < 0.01, *** P < 0.001, n.s. = not significant) **d.** Schematics of xenografting of SU-DIPGVI WT/PIEZO1 KO into immunocompromised mice followed by assessment of proliferation with Ki67+/HNA staining. **e.** Representative confocal images of WT and PIEZO1 KO brain slices explained in d. (white = Ki67, red = HNA, white arrows denote the overlapping cells with both Ki67 and HNA stained). **f.** Summary of tumor proliferation (g, %Ki67+) in WT and PIEZO1 KO samples. (n (number of independent biological samples) = 5, mean ± s.e.m., two-tailed t-test, * P < 0.05). **g.** Volcano plot of differentially expressed genes comparing WT and PIEZO1 KO MGG8 cells. Genes enriched in PIEZO1 KO relative to WT cells are plotted with positive log2 fold change. Conversely, genes that are depleted in PIEZO1 KO cells relative to WT are plotted as negative log2 fold change. Genes are colored according to log2 fold change and adjusted p-value thresholds (gray = non-significant by either metric, green = log2 fold change of 1 or greater, blue = adjusted p-value of 10e6 or greater, red = gene exceeds thresholds for fold change and p- value). **h.** Gene ontology analysis of genes that were significantly depleted in PIEZO1 KO cells relative to WT. Enriched programs all center around sterol biosynthesis and metabolism. Of note, FABP7 is amongst the most strongly down-regulated genes in PIEZO1 KO cells and is reported to function in the maintenance of both glioma stem cells and healthy OPCs^63^. **i.** Cartoon model depicting mechanism of NLGN3-induced CSPG4 shedding and functional consequences. Red circular sector = ADAM10; Green rectangle = shed NLGN3; blue = CSPG4 ectodomain; light blue = PIEZO1.

We then employed a less invasive approach of whole-cell recording electrophysiology with brief applications of hypertonic solution delivered through a glass pipette electrode to evoke mechanosensitive currents (Extended Data Fig. 3c). Studies from the Patapoutian lab^52^ and the Huang lab^53^ demonstrated that mechanical stress by osmotic gradient can ensure the preservation of cellular morphology, structural integrity, and membrane functionality. In line with previous work, we detected large inward currents following mechanical stimulation with hypertonic solution. Importantly, these responses were almost completely abolished in PIEZO1 CRISPR knockout cells, confirming that the current mainly arises from PIEZO1 conduction (Extended Data Fig. 3d and e). Notably, NLGN3 caused a dramatic change in the kinetics of the mechanical responses, notably slowing the inactivation phase (Extended Data Fig. 3f). This observation aligns well with the cell-attached recordings, and the data from CSPG4 shedding assays with modulation of mechanosensitive ion channels, to further confirm that NLGN3 is a regulator of PIEZO1 activity.

## Functional consequences in NLGN3-PIEZO1-mediated pathways

Having established that PIEZO1 is a primary actuator responsible for NLGN3-mediated shedding of CSPG4, we hypothesized that blocking the PIEZO1 response in glioma would mimic loss of microenvironmental NLGN3 and affect glioma growth in vivo^2,3^. To test this hypothesis, we knocked out PIEZO1 using CRISPR-Cas9 technology, and then orthotopically xenografted the PIEZO1 KO cells into pons region of mouse brains (Fig. 5d). Notably, after 4 weeks, mice bearing PIEZO1 KO glioma xenografts exhibited a significant reduction in glioma cell proliferation rate (Fig. 5e, f) compared to those bearing PIEZO1 WT glioma cells from the same patient. We further confirmed that administration of an ADAM10 inhibitor, which would block both NLGN3 secretion and NLGN3-mediated CSPG4 shedding, also resulted in a significant decrease in tumor burden and proliferation rate in patient-derived glioma xenografts, as expected^2^ (Extended Data Fig. 4). Taken together, these findings support the idea that that targeting the NLGN3 pathway, including PIEZO1 and ADAM10-mediated cleavage events, represent therapeutic strategies for gliomas.

To understand the transcriptomic consequences of PIEZO1 depletion, we next performed bulk RNA sequencing of a patient-derived glioblastoma cell culture models with and without PIEZO1 knockout via CRISPR-cas9 gene editing (Fig. 5h). We first confirmed by PCA analysis that WT and PIEZO1 KO samples clustered apart from one another. Next, we performed differential gene expression analysis and determined that upon PIEZO1 knockout (Fig. 5h), glioblastoma cells dramatically increase expression of several stress and apoptosis-related genes including CASP8, UBB and GSN. Intriguingly, down-regulated genes in PIEZO1-depleted cells were related to sterol metabolism (Fig. 5i), a program known to be highly active in glioblastoma cancer stem cells (CSCs)^54^. In line with this notion, several down-regulated genes are reported markers of glioblastoma CSCs including FABP7, TF and SLC2A3 (aka GLUT3), suggesting that knockdown of PIEZO1 may influence the maintenance of the cancer stem cell state (Fig. 5j)^55,56^.

## Blockade of CSPG4 shedding precludes NLGN3-induced changes in glioma

We next investigated whether the observed interaction between NLGN3 and CSPG4 is necessary for the NLGN3-induced changes in glioma, such as glioma proliferation and growth^1,2^. To test this hypothesis, we first tested the proliferation of glioma cells with and without NLGN3 treatment, using an EdU incorporation assay (Extended Data Fig. 5a). In agreement with prior studies^1–3^, we found that NLGN3 exposure increased proliferation of glioma cells (Extended Data Fig. 5b). Notably, this NLGN3-induced proliferation was attenuated by first pre-treating glioma cells with ADAM10 inhibitor, even in the presence of NLGN3 (Extended Data Fig. 5b), consistent with the hypothesized role for ADAM10-mediated CSPG4 cleavage in the NLGN3-CSPG4-PIEZO1 mechanism. Next, we investigated whether the knockout of the CSPG4 gene directly affects the growth of glioma cells using CSPG4 KO glioma cells and assessed cancer cell growth over time using the CellTiterGlo assay (Extended Data Fig. 5c). Indeed, we observed a significant reduction in the growth rate of CSPG4 KO glioma cells in neurosphere monoculture from day 3 onwards, indicating that CSPG4 may play a cell-intrinsic role in maintaining stem-like glioma neurospheres (Extended Data Fig. 5d).

## NLGN3-CSPG4 interaction impedes differentiation of OPCs

NLGN3 is expressed by and shed from both neurons and OPCs in the healthy brain^2^, but the role of NLGN3 in OPCs is unknown. Having confirmed that NLGN3 induces CSPG4 shedding from OPCs (Fig. 2-4), we investigated the activity of PIEZO1 in OPCs in response to NLGN3. Previous reports have demonstrated that PIEZO1 is a key mediator of OPC differentiation in the brain^6^. To confirm functional PIEZO1 expression, we conducted cell-attached patch clamp recordings in OPCs obtained from the brains of mouse pups (Extended Data Fig. 6a). We observed minuscule inward mechanosensitive currents from the mouse OPCs in response to negative pressure, which were notably augmented in the presence of NLGN3 (Extended Data Fig. 6b). Moreover, we found that the magnitude of these currents increased in response to the increasing magnitude of negative pressure (Extended Data Fig. 6c), akin to what we observed in PIEZO1 currents from glioma cells (Fig. 4). These data support the notion that PIEZO1 is indeed active in OPCs and raise the intriguing possibility that NLGN3-CSPG4 interactions may regulate OPC differentiation through PIEZO1 activity. We therefore investigated whether NLGN3 exposure influences OPC proliferation or differentiation *in vitro*. We first investigated whether NLGN3 can promote proliferation in OPCs, both in place of and in addition to the growth factor PDGFAA, a canonical OPC mitogen (Fig. 6a)^1,3^. In contrast to robust proliferation induced by PDGFAA, OPCs did not proliferate in response to NLGN3 exposure (Fig. 6b and c). PDGFAA growth factor withdrawal triggers oligodendrocyte differentiation which is characterized by dramatic and stereotypical morphological changes. We found that NLGN3 exposure impeded these morphological changes OPCs undergo to differentiate into oligodendrocytes following PDGFAA withdrawal (Fig. 6d and e). OPCs in medium containing PDGFAA typically extend single processes from opposite sides of their somas, while differentiating oligodendroglial cells become more complex and arborized (Fig. 6e and f)^57^. We reasoned that if these changes were due to NLGN3-mediated cleavage of CSPG4, then pre-treating OPCs with an ADAM10 inhibitor would render them insensitive to NLGN3 treatment and PDGFAA growth factor withdrawal would trigger their differentiation into oligodendrocytes. Indeed, blocking NLGN3-CSPG4 signaling via ADAM10 inhibition partially restored the morphological progression of OPC differentiation (Fig. 6g and h).

**Fig. 6.**
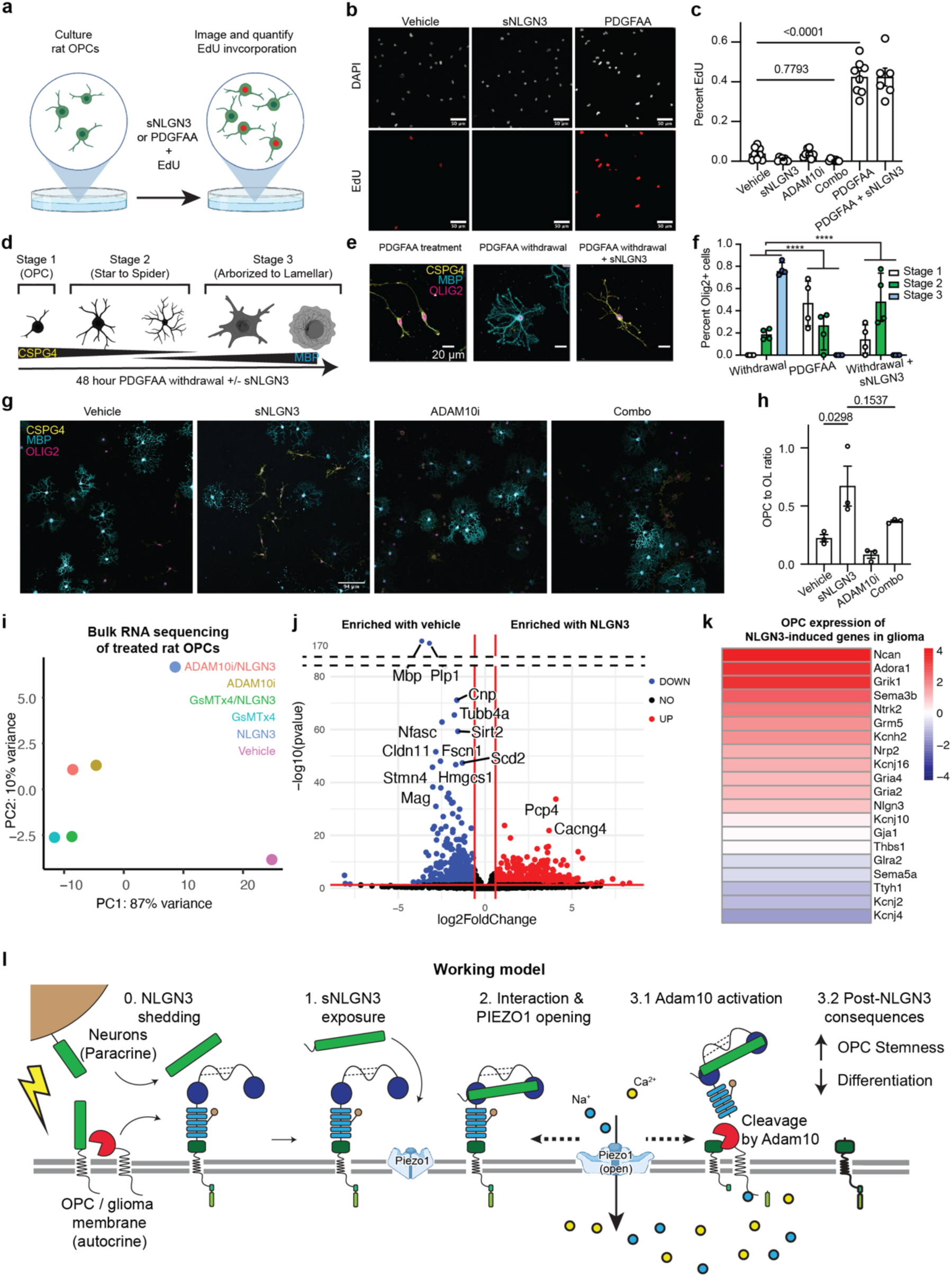
sNLGN3-induced CSPG4 shedding regulates OPC cell state. **a.** Workflow depicting EdU incorporation and detection in rat OPCs following treatment with NLGN3. **b.** Representative images of EdU incorporation in primary rat OPCs after 4-hour treatments. **c.** Quantification of cell proliferation depicted in a. (n (independent biological replicates) = 3) mean ± s.e.m., one-way ANOVA with Tukey’s multiple comparisons correction. **** P < 0.0001) **d.** Schematic depicting differentiation and morphological changes of OPCs upon mitogen withdrawal *in vitro*. **e.** Representative images of OPCs following 48-hour mitogen withdrawal in the presence or absence of supplemental sNLGN3. **f.** Quantification of cells at each morphological stage following 48-hours of treatment. ( n (number of independent trials) = 4, two-way ANOVA with multiple comparisons and Bonferroni correction, not significant) Note that PDGFAA treated and NLGN3-treated cells are not significantly different regarding progression to stage 3 morphology as assessed by two-way ANOVA with multiple comparisons and Bonferroni correction. **g.** Representative images of OPCs following 48-hour mitogen withdrawal in the presence or absence of supplemental sNLGN3 and ADAM10 inhibition. **h.** Quantification of OPC (CSPG4 positive) to OL (MBP positive) ratio in cells depicted in g (n (independent biological replicates) = 3) mean ± s.e.m., one-way ANOVA with Tukey’s multiple comparisons correction. p-values are written on the figures). **i.** PCA plot of bulk RNA-seq produced from rat OPCs that were cultured for 48 hours under PDGFAA withdrawal and supplementation with NLGN3 and/or inhibitors of CSPG4 shedding. Vehicle and NLGN3-treated cells are distinct in PC space from the inhibitor-treated samples. **j.** Volcano plot of differentially expressed genes comparing NLGN3-treated rat OPCs against PDGFAA withdrawal alone. Genes enriched in NLGN3-treated relative to WT cells are plotted with positive log2 fold change. Conversely, genes that are depleted in NLGN3-treated cells relative to WT are plotted as negative log2 fold change. Genes are colored according to log2 fold change and adjusted p-value thresholds (gray = non-significant by either metric, green = log2 fold change of 1 or greater, blue = adjusted p-value of 10e6 or greater, red = gene exceeds thresholds for fold change and p-value). **k.** Heatmap of NLGN3-induced genes from Venkatesh et al 2017. Primary rat OPCs were cultured for 48 hours without PDGFAA to initiate differentiation and NLGN3 was supplemented in half the cultures. Duplicate vehicle and NLGN3-treated samples were profiled by bulk RNA-seq. TPM values for each condition were averaged and then the log2 fold change was calculated between NLGN3-treated and vehicle control samples. These values were plotted such that zero appears as white on a symmetrical axis that ranges from -4 in blue and 4 in red. **l.** Model depicting CSPG4 shedding via PIEZO1 opening and ADAM10 activation. These events impede OPC differentiation in vitro upon growth factor withdrawal. Red circular sector = ADAM10; Green rectangle = shed NLGN3; blue = CSPG4 ectodomain; light blue = PIEZO1.

To corroborate these morphological changes at the molecular level, we next assessed transcriptomic changes via bulk RNA sequencing of rat OPCs undergoing spontaneous differentiation in the presence or absence of NLGN3 and modulators of CSPG4 shedding. We determined by PCA analysis that OPCs treated with NLGN3 clustered apart from cells exposed to vehicle or pre-treated with inhibitors of CSPG4 shedding prior to NLGN3 exposure (Fig. 6i). Furthermore, our differential gene expression analysis comparing NLGN3-treated OPCs and vehicle-treated OPCs clearly demonstrates an enrichment of genes associated with stemness and synaptic communication (*Pdgfra, Cacng4,* and *Kcnip3*) and a suppression of genes related to differentiation (*Mbp, Plp, Enpp6*) (Fig. 6j). Of note, we found that NLGN3 treatment of OPCs resulted in upregulation of several genes that also increased in expression when human glioma cells were treated with NLGN3 in a previous study demonstrating the functional similarities of Nlgn3 in OPCs and glioma cells (Fig. 6k)^2^. Finally, using RT-qPCR analyses in OPCs, we verified that NLGN3 treatment prevented the expression of myelin-specific transcripts Plp and Mbp and maintained the expression of the differentiation inhibitor Sox11 relative to OPCs exposed only to vehicle control following growth factor withdrawal (Extended Data Fig. 6d)^58^.

Previous in vivo genetic mouse modeling experiments demonstrated that NLGN3 is chiefly shed from OPCs, implicating that OPC-derived NLGN3 shedding may play a role in oligodendroglial biology. The in vitro studies described above implicate NLGN3 signaling in maintenance of a self-renewing, undifferentiated state in OPCs, raising the possibility that NLGN3 shed from OPCs may contribute to OPC stemness and population maintenance. We therefore tested the effects of *Nlgn3* loss on OPC population homeostasis using a conditional knockout model, PDGFRa::Cre^ERT^;Nlgn3^fl/fl^, where Nlgn3 is specifically knocked out from OPCs upon tamoxifen administration (Extended Data Fig. 6e). Remarkably, acute deletion of NLGN3 led to an approximately 40% reduction in the number of OPCs in the corpus callosum (Extended Data Fig. 6f, g; Fig. 6l).

## Discussion

Neuronal activity regulates the behavior of healthy glial and malignant glioma cells. In the healthy nervous system, activity causes adaptive oligodendrocyte precursor cell proliferation, generation of new oligodendrocytes, and myelination and this plasticity of the oligodendroglial lineage tunes neural circuits to support healthy brain functions like learning and memory (For review, please see ^10^). Malignant gliomas – many of which arise from OPCs and which contain important cellular subpopulations that closely resemble OPCs – are similarly regulated by neuronal activity and proliferate in response to activity-regulated paracrine factors and direct neuron-to-glioma synaptic communication. Membrane depolarization alone is sufficient to promote glioma cell proliferation^16,22^. One key paracrine factor released as a result of neuronal activity is Neuroligin-3 (NLGN3), originally known for its role as a synaptic adhesion molecule. However, the precise mechanisms mediating NLGN3 interactions with glioma surface signaling proteins remained elusive.

Here, we uncovered a mechanism by which NLGN3 interacts with CSPG4, thereby altering membrane tension and activating the mechanosensitive ion channel PIEZO1 with consequent membrane depolarization and depolarization-induced activity of the ADAM10 protease. This NLGN3-CSPG4-PIEZO1 axis represents a novel convergence point between neuronal signaling^4,10,59,60^ and mechanotransduction pathways in glioma cells^53,61^. This interaction underscores cancer’s adaptability in harnessing both biochemical and mechanical cues from their microenvironment to sustain growth. Beyond its implications for glioma biology, these findings shed new light on roles for NLGN3 in the regulation of OPC states, elucidating a newly appreciated role for activity-regulated maintenance of homeostatic OPC population dynamics through the NLGN3-CSPG4-PIEZO1 axis. NLGN3-CSPG4-PIEZO1 signaling thus joins an ever-growing list of mechanistic parallels between neuron-glial and neuron-glioma interactions, promoting a stem-like, OPC state in both healthy and malignant glial cells. In healthy OPCs, this stem-like state prevents differentiation while in glioma it drives proliferation.

In summary, the findings presented here underscore a critical role for NLGN3 in the interplay between neuronal activity and glioma progression, unveiling a newly appreciated mechanistic pathway by which gliomas exploit neuronal signals for growth. The identification of PIEZO1 as a key mediator in this process suggests new therapeutic opportunities, highlighting the potential for targeting mechanosensitive pathways in glioma and possibly in other cancers.

## MATERIALS AND METHODS

### Mice and housing conditions

All animal experiments were conducted in accordance with protocols approved by the Stanford University Institutional Animal Care and Use Committee (IACUC) and performed in accordance with institutional guidelines. Animals were housed according to standard guidelines with unlimited access to water and food, under a 12 hour light : 12 hour dark cycle. For brain tumor xenograft experiments, the IACUC has a limit on indications of morbidity (as opposed to tumor volume). Under no circumstances did any of the experiments exceed the limits indicated and mice were immediately euthanized if they exhibited signs of neurological morbidity or if they lost 15% or more of their initial body weight.

### Recombinant NLGN3 purification

DNA sequence encoding the signal peptide and ectodomain of human NLGN3 (Sino Biological NLGN3=HG11160-UT) and subsequently cloned into a mammalian cell expression vector (pD649) with AviTag and 6xHis Tag at the C-terminus. These plasmids were transfected into Expi293 cell culture, followed by induction after 18-24 hours of transfection. After 72-96 hours of incubation, media were harvested and filtered via 0.22-micron Steriflip^TM^. Recombinant proteins were then captured via cobalt affinity chromatography (HisPur Cobalt Resin, Thermo 89964), followed by washing with 20 column volume of wash buffer (100 mM NaCl, 20 mM HEPES pH 7.5, 20 mM imidazole) and elution with with 5 CV of elution buffer (100 mM NaCl, 20 mM HEPES pH 7.5, 250 mM imidazole). Eluents were then subjected to gel filtration chromatography in the final buffer (100 mM NaCl, 20 mM HEPES pH 7.5) and thenconcentrated to 3-6 mg/ml for experiments.

### NLGN3 affinity capture assay

30 million SU-DIPG13pons or MGG8 cells were cultured as neurospheres and pelleted without dissociation by spinning (200g for 3 min, room temperature). Cells were washed in HBSS (w/o calcium and magnesium), pelleted once more, and then snap-frozen in dry ice/ethanol and kept at -80°C to facilitate downstream cell lysis and membrane isolation. Pellets were thawed on ice, then resuspended in 5 mL ice-cold lysis buffer supplemented with protease inhibitors (10 mM HEPES pH 7.5, 10 mM MgCl_2_, 20 mM KCl + 1x Halt^TM^ Protease inhibitor cocktail (Thermo Fisher, 78429)). Samples were then spun at 20,000g for 20 min at 4C and supernatants were assessed for protein content by Bradford assay. Pellets were resuspended in 1 mL of chilled lysis buffer with protease inhibitors and homogenized with 20 strokes of a 2mL glass dounce. Lysates were diluted to 4 mL in lysis buffer and spun at 20,000 G for 20 min at 4C. This homogenization procedure was repeated three times. Following the last spin, pellets were resuspended in 2.5 mL of salt wash buffer (lysis buffer + 150 mM NaCl) with protease inhibitors. Homogenization with the dounce was repeated in this salt wash buffer until no protein was detected via Bradford assay in the cleared supernatant after spinning. We then considered pellets to be purified membrane fractions and solubilized these in (1% L-MNG, 150mM NaCl, 20mM HEPES, 10% glycerol, supplemented with 1x Halt protease inhibitor cocktail) for incubation with NLGN3 affinity resin. Separately, we biotinylated our homemade NLGN3 ectodomains using the BirA biotin-protein ligase standard reaction kit (Avidity, EC 6.3.4.15). Proteins were then purified of excess biotin and concentrated via a 30K MWCO Amicon spin filter (MilliporeSigma UFC503096). Biotinylated NLGN3 was then immobilized as bait on streptavidin resin (Pierce ProFound™ Biotinylated-Protein Interaction Pull Down Kit, Thermo 21115). Solubilized membrane proteins were incubated with NLGN3-resin overnight with rotation at 4C, washed three times with supplied acetate buffer supplemented with 150 mM NaCl and then eluted by incubating in LDS sample buffer + BME at 42F for 15 minutes. Proteins were resolved on a 3-8% tris-acetate gel run at 120V for 15 min before slices near the top of the gel were excised for evaluation by mass spectrometry.

For calculating the enrichment index, the amount of proteins identified by mass spectrometry was normalized by the amount of actin in each dataset. After normalization, the amounts of proteins were compared with and without NLGN3 affinity capture. The enrichment index was calculated by dividing the amount of each protein with NLGN3 capture by the amount of the same protein identified without NLGN3 capture.

### Western blots from human CSF

Human CSF samples were procured under the IRB14511, from the Bass Center Leukemia Tissue Bank (for non-glioma, acute lymphocytic leukemia patient CSF samples) and from the GD2-CAR-T cell clinical trial (for glioma patient CSF samples, from Dr. Sneha Ramakrishna). Procured samples were concentrated (Amicon 50K filter) 20-fold.

For western blot analysis, samples were prepared by adding LDS buffer and β-mercaptoethanol, heated, and loaded onto Tris-acetate gels. After electrophoresis and transfer onto PVDF membranes, the membranes were blocked and incubated with primary antibodies against the NLGN3 ectodomain (Abcam #90080) N-terminal regions of CSPG4 (Cell Signaling Tech #52635). Following incubation with secondary antibody and washing, the membranes were developed using chemiluminescent substrate (Thermo Scientific #34580).

### NLGN3 coupling assay using Fluorescence Size Exclusion Chromatography

For initial coupling analysis, purified NLGN3 (320µg/ml) was mixed with 6 mg/ml of Chondroitin Sulfate (CS, Sigma Aldrich) or 400 µg/ml of heparin (heparan sulfate, US Biological). Coupling was assessed by the left shift in Fluorescence Size Exclusion Chromatography (FSEC)^64^, using tryptophan fluorescence for detecting NLGN3 signal. For quantification, we mixed the same amount of NLGN3 with 0, 0.75, 1.5, 3, and 6 mg/ml of CS and quantified the left shift of the NLGN3 peak in FSEC. Experiments were conducted three times, with independently purified batches of NLGN3.

### Glioma Cell Culture

DIPG or GBM neurospheres (SU-DIPG 6, SU-DIPG13pons, SU-DIPG21, MGG8) were cultured in defined medium composed of a 1:1 mixture of Neurobasal and DMEM F12, supplemented with Glutamax, penicillin-streptomycin, MEM non-essential amino acids (NEAA), sodium pyruvate, HEPES, B-27 supplement (without vitamin A), heparin, and growth factors (EGF, FGF, PDGF-AA, PDGF-BB). The cultures were maintained at 37°C with 5% CO2.

### Primary Oligodendrocyte Precursor Cell Culture

Primary oligodendrocyte precursors were purified by immunopanning from P5-P7 Sprague Dawley rat and P6-P7 transgenic mouse brains as previously described^65^. They were seeded at 150,000–250,000 cells per 10 cm dish, recovered for 4 days, then trypsinized for assays.

Plasticware was coated with 0.01 mg/ml poly-D-lysine (PDL, Sigma P6407); glass coverslips with 0.01 mg/ml PDL in 150 mM boric acid, pH 8.4.

For proliferation, cells were cultured in serum-free DMEM-SATO with 4.2 μg/ml forskolin (Sigma-Aldrich, F6886), 10 ng/ml PDGF (Peprotech, 100-13A), 10 ng/ml CNTF (Peprotech, 450-02), and 1 ng/ml NT-3 (Peprotech, 450-03) at 37°C, 10% CO2. For differentiation, DMEM-SATO was supplemented with 4.2 μg/ml forskolin, 10 ng/ml CNTF, 40 ng/ml thyroid hormone (T3; Sigma-Aldrich, T6397), and 1× N21-MAX (R&D Systems AR008).

### CSPG4 shedding assay

For glioma cells, neurospheres were pelleted, washed, and dissociated into single cells using 5 mM EDTA in HBSS (without calcium and magnesium). The dissociation process involved gentle trituration until complete dissociation into single cells was achieved. Dissociated cells were resuspended in HBSS with calcium and magnesium, counted, and plated into wells of a 12-well plate.

Cells were treated with 5 µM ADAM10 inhibitor (GI254023X), 2.5 mM 4-AP, 2 µM cytochalasin D, 10 µM Yoda-1, 10 µM gadolinium chloride, 2.5 μM GsMTx4, or DMSO control for 15 minutes. Recombinant human NLGN3 ectodomain or matched vehicle was added to the wells, and cells were incubated for 1 hour. Following incubation, the conditioned medium was collected by pipetting into Eppendorf tubes. The tubes were centrifuged to pellet any cell debris, and the cleared supernatant was collected. The supernatant was concentrated using 30K MWCO Amicon spin filters until the volume was reduced to 40 uL.

For western blot analysis, samples were prepared by adding LDS buffer and β-mercaptoethanol, heated, and loaded onto Tris-acetate gels. After electrophoresis and transfer onto PVDF membranes, the membranes were blocked and incubated with primary antibodies against the C-terminal and N-terminal regions of CSPG4. Following incubation with secondary antibody and washing, the membranes were developed using chemiluminescent substrate.

Pixel intensities of CSPG4 protein bands were quantified using FIJI software and reported as fold changes relative to vehicle control samples. Longer exposures were obtained for illustrative purposes.

This protocol was also adapted for the shedding assay using mouse optic-pathway glioma cells, normal mouse, and rat OPCs. OPCs were isolated and plated, and the assay was performed following a similar procedure. Experiments were replicated at least three times to ensure reproducibility and reliability of results.

### Live Cell Membrane Tension Measurement

For the measurement of membrane tension, primary cultured cells were prepared on coverslips coated with Poly-L-Lysine. The staining solution was prepared by mixing 1µM membrane tension sensor Flipper-TR (Spirochrome) with the imaging solution (150 mM NaCl, 10 mM HEPES pH 7.3, 10 mM glucose, 2 mM CaCl2, 2mM MgCl2, 4 mM KCl), and the staining solution was replaced with the staining solution and was incubated at 37°C, 5% CO2 for 30 minutes before imaging. No impact on cell viability has been observed after 2-3 days of incubation. For the NLGN3 incubation, 400 nM NLGN3 was added to the staining solution, and for the negative control, the same concentration of bovine serum albumin or mouse IgG was added.

Cells are imaged with standard FLIM microscopes using a 488 nm pulsed laser for excitation and collecting photons through a 600/50 nm bandpass filter.. To extract lifetime information, the photon histograms from ROI or single pixels (accumulate sufficient counts to ensure good statistics) were fitted with a double exponential, and two decay times, tau1 and tau2 were extracted. The longest lifetime with the higher fit amplitude tau1 was used to report membrane tension and varies between 2.8 and 7.0 ns.

### Electrophysiology

For whole-cell recording experiments, primary cultured cells were recorded using the methods described previously^64^. Briefly, cells were placed in an extracellular medium (150 mM NaCl, 4 mM KCl, 2mM CaCl2, 2 mM MgCl2, 10 mM HEPES pH 7.3 – 7.4 and 10 mM glucose, osmolarity 335-345). A borosilicate patch pipette (Harvard Apparatus) with resistance of 5 – 6 MΩ was filled with intracellular medium (140 mM potassium-gluconate, 10 mM EGTA, 2 mM MgCl2 and 10 mM HEPES pH 7.2). Signals were amplified and digitized using the Multiclamp 700B and DigiData1400 (Molecular Devices), and Leica DM LFSA microscope were used to visualize the cells. Whole-cell voltage-clamp experiments were conducted at a membrane potential of −70 mV. For the optogenetics experiment, 1.0 mW/mm^2^ light of 460 nm was delivered for 1 second, and the resulting photocurrent was measured. For mechanosensitive current recording experiment, hypertonic solution (extracellular medium + 300 mM sucrose) or isotonic solution (extracellular medium) was used for mechanosensitive stimuli. The hypertonic high sucrose jet stream was used to evoke mechanosensitive currents via osmotic pressure-induced stretch of the cell membrane as previously described^53^.

For cell-attached PIEZO1 recording experiments, currents were recorded using the methods described previously^52^ external solution used to zero the membrane potential consisted of (in mM) 140 KCl, 1 MgCl_2_, 10 glucose, and 10 HEPES (pH 7.3 with KOH). Recording pipettes were of 3–6 MΩ resistance when filled with standard solution composed of (in mM) 130 NaCl, 5 KCl, 1 CaCl_2_, 1 MgCl_2_, 10 TEA-Cl, and 10 HEPES (pH 7.3 with NaOH). When specified, pipette solution was supplemented with 30 μM Yoda1, 10 μM GsMTx4, or 500 nM NLGN3. Membrane patches were stimulated with 500-ms or 1 second negative pressure pulses through the recording electrode using Clampex controlled pressure clamp HSPC-1 device (ALA-Scientific, Farmingdale NY). Consecutive sweeps with pressure stimulation ranging from 0 to −60 mm Hg (Δ-10 mm Hg) were applied every 30s.

All experiments were performed at the room temperature. Data were analyzed using pClamp10 software (Molecular Devices). Analysis of mechanically activated currents by fold change was performed using peak currents from the onset of the pressure step to 1.0 s to account for any late or secondary currents.

### Visualization of PIEZO1 and CSPG4 expression from single cell RNA-seq of high-grade gliomas

Gene expression matrices corresponding to Filbin et al., 2018^66^ and Neftel et al., 2019^67^ were downloaded from GEO and first analyzed via the Seurat software package to identify and remove normal microglia and oligodendrocytes from the datasets^68^. Cells were clustered according to the standard Seurat workflow and scored for microglia and oligodendrocyte gene expression programs. Both cell types form clear and discrete clusters that are removed by sub- setting the Seurat object for further analysis. Remaining cells were then scored according to gene expression modules that were identified in each paper (stem-like, AC-like, OC-like, OPC-like, MES-like, NPC-like) then visualized by ggplot2 according to these scores. The color of each cell corresponds to a scaled expression value for PIEZO1 (top row) or CSPG4 (bottom row). Further, cells were first ranked according to PIEZO1 expression before plotting such that cells with the highest expression were plotted last.

### CRISPR knockout of CSPG4 from high-grade glioma cells

sgRNA oligos targeting human CSPG4 were designed using the CRISPOR pipeline (http://crispor.tefor.net/crispor.py). Oligos were annealed and cloned into pL-CRISPR.EFS.GFP (Addgene # 57818) using BsmbI according to the Juong lab protocol available on Addgene. Plasmids containing CSPG4-targeting sgRNA were verified by sanger sequencing. This plasmid was transfected with helper plasmids in 293T cells to produce replication deficient lentivirus. An empty pL-CRISPR.EFS.GFP plasmid was used to produce non-targeting lentivirus for controls. Collected viral supernatant was spun at 500 G for 5 min, then filtered through a 0.45 micron filter before being concentrated using the Takara LentiX Concentrator (Takara, 631231) reagent according to manufacturer’s instructions. Concentrated virus titer was approximated using the Takara Lenti-X GoStix Plus (631280) system according to manufacturer’s instructions. Glioma neurospheres were dissociated to single cells and 1.5 million cells were plated in 2 mL of complete medium in one well of a six-well plate and allowed to recover for 2 hours in the incubator before being infected with control or CSPG4 targeting lentivirus. After 96 hours, CSPG4 KO cells were isolated via FACS. Following FACS isolation, cells were returned to regular medium for propagation.

### FACS of CSPG4 KO high-grade glioma cells

96-hours after infection with non-targeting or CSPG4-targeting sgRNA and Cas9, high-grade glioma neurospheres were dissociated to single cell suspensions and stained with an alexa fluor 647-labeled anti-CSPG4 antibody (BD Biosciences, 562414) or an isotype-matched control antibody. CSPG4-poisitive cells were sorted into a 6-well TC plate containing complete medium using a BD Aria and following BL2+ containment practices.

### CRISPR knockout of PIEZO1 from high-grade glioma cells

DNA plasmid encoding a gRNA targeting human PIEZO1 was purchased from Genscript (PIEZO1 CRISPR guide RNA 3_pLentiCRISPR v2, CATCCCCAACGCCATCCGGC) and transfected with helper plasmids in 293T cells to produce replication deficient lentivirus. An empty pLentiCRISPR v2 plasmid was used to produce non-targeting lentivirus for controls. Collected viral supernatant was spun at 500 G for 5 min, then filtered through a 0.45 micron filter before being concentrated using the Takara LentiX Concentrator (Takara, 631231) reagent according to manufacturer’s instructions. Concentrated virus titer was approximated using the Takara Lenti-X GoStix Plus (631280) system according to manufacturer’s instructions. Glioma neurospheres were dissociated to single cells and 1.5 million cells were plated in 2 mL of complete medium in one well of a six-well plate and allowed to recover for 2 hours in the incubator before being infected with control or PIEZO1 targeting lentivirus. After 48 hours, 0.5 – 1 ug/mL of puromycin was added to culture medium to select for successfully infected cells. Control cells without puromycin were monitored to ensure loss of viability. Depletion of PIEZO1 protein levels from cells was confirmed via western blot. Following puromycin selection, cells were returned to regular medium for propagation.

### Bulk RNA of glioma cells and rat OPCs

Glioma neurospheres were harvested and dissociated to a single cell suspension using ACCUTASE (Stem Cell Technologies, 07920). Cells were counted and 400,000 were pelleted. Cell pellets were lysed in 350 uL microliters of buffer RLT (Qiagen, 74134) and total RNA isolated following manufacturer’s instructions for the Rneasy Mini kit. 200 nanograms of total RNA for each sample was used as input for reverse transcription with the Template Switching RT Enzyme Mix (NEB, M0466L). The RT primer, template switching oligo (TSO) and cDNA amplification primer (ISPCR) were sourced from the Smart-seq2 protocol^69^. Oligo-dT30VN (5′–AAGCAGTGGTATCAACGCAGAGTACT30VN-3′) TSO (5′-AAGCAGTGGTATCAACGCAGAGTACATrGrG+G-3′) ISPCR (5′-AAGCAGTGGTATCAACGCAGAGT-3

cDNA was then amplified with ISPCR primer and 2x KAPA HiFi readymix (Roche, KK2102). Amplified cDNA was then cleaned by 2x SPRI bead purification and measured by the Qubit 2.0 fluorometer. Samples were normalized to 0.2 ng/uL and a total of 0.8 ng of cDNA was used as input for tagmentation reaction according to manufacturer’s instructions (Illumina, FC-131- 1096). Tagmented DNA was then combined with dual unique indexes (Illumina, 20027213) and amplified via PCR according to manufacturer’s instructions. Final libraries were cleaned by 0.6x SPRI and analyzed using the high-sensitivity DNA bioanalyzer kit (Agilent, 50674626). Bioanalyzer traces were used to quantify total DNA between 200 and 1000 bp before libraries were combined such that the final pool was at 2 nanomolar. The pool was further diluted to 650 picomolar and loaded into a P2 kit (Illumina, 20046812) for sequencing on the Illumina NextSeq 2000 (2 x 100 bp reads, 2 x 10 bp indices). Rat OPCs were processed similarly with the following modifications. 85,000 rat OPCs were plated onto PDL-coated wells of a 12-well plate and cultured for 48 hours in the absence of PDGFAA. Instead, OPCs were treated with 400 nanomolar NLGN3, 5 uM ADAM10i, 5 uM GsMTX4, and combinations thereof. Following 48- hour incubation. Medium was aspirated and 350 uL of buffer RLT was added directly to the plate to lyse cells.

For the alignment and quantification of bulk RNA sequencing from glioma cells and rat OPCs, demultiplexed FASTQ files were first pre-processed for quality via fastp (https://academic.oup.com/bioinformatics/article/34/17/i884/5093234) then aligned to the human or rat transcriptome via Salmon version 0.12.0^70^. Quantifications from Salmon were then passed through Tx-import and summarized at the gene level according to the standard DeSeq2 workflow(https://www.bioconductor.org/packages/release/bioc/vignettes/DESeq2/inst/doc/DESeq2.html#transcript-abundance-files-and-tximport-tximeta).

### Single cell RNAseq

#### Differential gene expression analysis of PIEZO1 WT and KO glioma cells and rat OPCs

Differential gene expression analysis was performed using the DESeq2 package with references set for each experiment separately (PIEZO1 WT glioma cells and vehicle-treated rat OPCs). The results table was downloaded and used to construct a volcano plot of significantly up- and down- regulated genes with the EnhancedVolcano R package (https://bioconductor.org/packages/release/bioc/html/EnhancedVolcano.html).

#### Scoring of rat OPCs for NLGN3-induced genes from Venkatesh et al., 2017

The log2 transformed TPM expression values for genes that were significantly up-regulated in pcGBM2 following NLGN3 treatment were extracted from the rat OPC samples treated with NLGN3 or vehicle and plotted as a heatmap using the pHeatmap R package (https://www.rdocumentation.org/packages/pheatmap/versions/1.0.12/topics/pheatmap).

#### Gene ontology analysis of PIEZO1 WT and KO glioma cells

Differentially expressed genes identified by DESeq2 were used as input for gene ontology analysis with the enrichGO R package (https://www.rdocumentation.org/packages/clusterProfiler/versions/3.0.4/topics/enrichGO).

#### EdU incorporation assay

10,000 optic glioma cells were seeded to fibronectin (10 μg ml^-1^)-coated 8-chamber slides (Thermo Fisher Scientific, 154534PK), and grown in NSC medium without N2, FGF and EGF for 48 h, supplemented with vehicle (150mM NaCl, 50 mM HEPES), recombinant human NLGN3 ectodomain at 100 nM (produced in house, see methods) at the time of plating and again 24-hours following plating. EdU proliferation assay was performed using the Click-iT EdU Alexa Fluor 594 imaging kit (Thermo Fisher Scientific C10339) according to the manufacturer’s instructions. EdU (10 uM) was added 12 hours prior to fixing cells in 4% paraformaldehyde. Images were taken using a Zeiss Axio Imager M2 and the Stereo Investigator software (mbf bioscience v2019). Cell proliferation was determined by dividing the number of EdU-positive cells by the number of DAPI-positive cells using Cell Profiler (v3.1.9). Each point in the graph represents the average value of 2-independent wells of cells and 3 images per well. Rat OPCs were initially plated and expanded in PDGFAA-containing medium. Cells were lifted in the afternoon by trypsin and 10,000 cells were replated in 100 ul bubbles on dried PDL-coated glass coverslips. After 30 minutes in the incubator to adhere, an additional 400 uL of base medium was added to cells. Cells were left overnight to incubate at 37C at 10% CO2. The next morning, cells were pulsed with 100 uL of base medium containing EdU to achieve a final concentration of 10 micromolar. The added 100 uL also contained NLGN3, ADAM10 inhibitor and/or PDGFAA. Cells were returned to the incubator for 4-hours before being fixed in 4% paraformaldehyde. Cells were stained using the Click-iT™ EdU Alexa Fluor™ 594 Imaging Kit and subsequently stained with DAPI according to manufacturer’s instructions. 2-D snaps of EdU and DAPI were taken with an LSM700 by a blinded investigator. Furthermore, fields of view were chosen with the DAPI channel to avoid bias in selecting regions for counting.

### High-grade glioma xenografts of wild-type and PIEZO1 knockout cells

For all xenograft studies, NSG mice (NOD-SCID-IL2R gamma chain-deficient, The Jackson Laboratory) were used. Male and female mice were used equally. For all experiments, SU- DIPG6, SU-DIPG13pons or pcGBM2 neurospheres were dissociated using 1x Accustase. Cells were counted with Trypan Blue. Cells were pelleted and resuspended in a volume such that 100,000 cells were suspended in 1 uL of HBSS and placed on ice. Animals at postnatal day (P) 28 were anaesthetized with 1–4% isoflurane and placed in a stereotactic apparatus. The cranium was exposed via midline incision under aseptic conditions. Approximately 300,000 cells in 3 µl sterile HBSS were stereotactically implanted into cortex (0.5 mm lateral to midline, 1 mm anterior to bregma, −1.0 mm deep to cranial surface) or the pons (1.0 mm lateral to midline, −0.8 mm posterior to lambda, −5.0 mm deep to cranial surface) through a burr hole, using a digital pump at infusion rate of 0.4 µl min−1 and 26-gauge Hamilton syringe. At the completion of infusion, the syringe needle was allowed to remain in place for a minimum of 2 min, then manually withdrawn at a rate of 0.875 mm min−1 to minimize backflow of the injected cell suspension.

### Immunohistochemistry of PIEZO1 KO/WT SU-DIPG6 mice

Mice were anaesthetized with intraperitoneal avertin (tribromoethanol), then transcardially perfused with 20 ml of PBS. Brains were fixed in 4% PFA overnight at 4 °C, then transferred to 30% sucrose for cryoprotection. Brains were then embedded in Tissue-Tek O.C.T. (Sakura) and sectioned in the coronal plane at 40 μm using a sliding microtome (AO 860, American Optical). For immunohistochemistry, coronal or sagittal sections were incubated in blocking solution (3% normal donkey serum, 0.3% Triton X-100 in TBS) at room temperature for 30 min. Rabbit anti- Ki67 antibody (1:500, Abcam, ab15580) was diluted in antibody diluent solution (1% normal donkey serum in 0.3% Triton X-100 in TBS) and incubated overnight at 4 °C. Sections were then rinsed once with TBS, before an incubation with DAPI (1 µg ml^−1^ in TBS, Thermo Fisher Scientific) and then another rinse with TBS. Slices were incubated in secondary antibody solution; Alexa 594 donkey anti-rabbit IgG was used at 1:500 (Jackson Immuno Research) in antibody diluent at 4 °C overnight. Sections were washed three times with TBS and mounted with ProLong Gold Mounting medium (Life Technologies). Confocal images were acquired on either a Zeiss Airyscan1 800 or Zeiss Airyscan LSM980 using Zen 2011 v8.1. Proliferation index was determined by quantifying the fraction of Ki67-labelled cells/human nuclear antigen (HNA)-labelled cells using confocal microscopy at 20x magnification. Quantification of images was performed by a blinded investigator.

### Confocal Imaging of SU-DIPG13pons mice

For 3D reconstruction, SU-DIPG13 pons xenografted mice were treated in the same manner as for immunohistochemistry of SU-DIPG6 mice, using a 1:500 dilution of rabbit PIEZO1 antibody (ProteinTech 15939-1-AP) and chicken anti-GFP antibody (Abcam ab13970). This was followed by the application of anti-chicken 488 (Jackson ImmunoResearch 703-545-155) and anti-rabbit 594 secondary antibodies (Jackson ImmunoResearch 111-585-144). Images were processed using IMARIS (v10.0.0) software.

### OPC differentiation assay

Rat OPCs were initially plated and expanded in PDGFAA-containing medium as described^65^. Cells were lifted by trypsin and 10,000 cells were replated in 100 ul drops on dried PDL-coated glass coverslips. After 20 minutes in the incubator to adhere, an additional 400 uL of base medium was added to cells. The medium was then supplemented with 5 µM DMSO as a vehicle, 20 ng/mL PDGFAA, 400 nM NLGN3, 5 µM ADAM10 inhibitor, 5 µM GsMTx4, or combinations thereof as specified. Cells were incubated at 37 C and 10% CO2 for 24-hours then supplemented with additional NLGN3 before being fixed with 4% paraformaldehyde at 48- hours. Cells were blocked and permeabilized in 3% normal donkey serum and 0.5% triton-x 100 for 30 minutes at room temperature before incubation with 1:1000 dilutions of anti-MBP (Abcam ab7349), CSPG4 (gift from Dr. Jacqueline Trotter), and OLIG2 (R&D AF2418) overnight at 4C in 1% normal donkey serum and 0.3% triton-x 100. Slips were washed 3 times in TBS for 5 minutes at room temperature before being incubated with 1:500 of fluorescent secondary antibodies (Jackson ImmunoResearch) for 1 hour at room temperature. Slips were washed 3 more times in TBS for 5 minutes each at room temperature, let to dry completely in the dark, then mounted onto microscope slides for imaging with Prolong Gold (P36930).

### Quantification of OPC morphological stage

Cells were examined with an LSM700 confocal microscope using a 20X objective. In all treatment groups, fields of view were chosen using the DAPI channel to avoid any bias. Fields were chosen haphazardly with a goal of capturing evenly spaced nuclei. Once chosen, a 2-D snap of the field was taken with all four channels. Two snaps were taken per coverslip and two coverslips were processed per experiment. This experiment was then replicated. The FIJI Cell Counter plugin was then used to facilitate qualitative assignment of cells to various buckets based on morphology. First, all DAPI nuclei in the field of view were counted. Next OLIG2-positive, DAPI+ nuclei were counted. Finally, cells were classified as exhibiting OPC-like, star and spider morphology using the CSPG4 channel. The MBP channel was used to classify cells as belonging to the arborized to lamellar category. These data were then reported as total number of cells in each “stage” of differentiation. This way of displaying the data allows one to appreciate that fewer cells are left in the PDFGAA withdrawal condition, indicating cell death. Conversely, the addition of NLGN3 appears to preserve cell numbers, while the continuous presence of PDGFAA induces proliferation as expected.

### Quantification of CSPG4:MBP ratio in primary rat OPCs

Rat OPCs from the differentiation assay were also evaluated for expression of oligodendrocyte lineage markers CSPG4 (OPCs) and MBP (oligodendrocytes). For each experiment, treatment conditions were replicated across 3 coverslips. A total of eight z-stacks were obtained for each coverslip that were subsequently converted to maximum intensity projections. Total numbers of OLIG2-positive nuclei, CSPG4-positive cells and MBP-positive cells were counted per field and reported as the ratio of CSPG4:MBP positive cells.

### qRT-PCR of rat OPCs undergoing spontaneous differentiation

Rat OPCs used in gene expression studies were identically manipulated as above. Cells were directly harvested from coverslips in 24-well plates. Cells were first differentiated as described above in the presence or absence of NLGN3 and then washed with HBSS before being lysed in buffer RLT from the Qiagen RNeasy mini kit. Total RNA was isolated and 100 ng from each experimental condition added to reverse transcription reaction using the New England Biolabs Protoscript II reverse transcriptase enzyme. Equal volumes of cDNA were added to each qRT-PCR reaction. Expression primers were previously validated ^71^. Fold change in gene expression was calculated using the delta-delta Ct method relative to GAPDH expression.

## Reporting Summary

Further information on research design is available in the Nature Research Reporting Summary linked to this paper.

## Data availability

All research materials are available upon request. All data reported in this paper will be shared by the corresponding authors upon request. Any additional information required to reanalyze the data reported in this paper is available from the corresponding authors upon request.

**Supplementary Information** is linked to the online version of the paper

## Acknowledgments

We are grateful to Sigrid Knemeyer at SciStories for illustrations. We acknowledge funding from the the National Institute of Neurological Disorders and Stroke (R01NS092597 to M.M.), NIH Director’s Pioneer Award (DP1NS111132 to M.M.), National Cancer Institute (P50CA165962, R01CA258384, U19CA264504 to M.M.), Gatsby Charitable Foundation (Gatsby Initiative in Brain Development and Psychiatry, to M.M.), Oscar’s Kids Foundation (to M.M.), McKenna Claire Foundation (to M.M.), Kyle O’Connell Foundation (to M.M.), Yuvaan Tiwari Foundation (to M.M.), Virginia and D.K. Ludwig Fund for Cancer Research (to M.M.), Waxman Family Research Fund (to M.M.), Will Irwin Research Fund of the Pediatric Cancer Research Foundation (to M.M.), Austin Strong Foundation (M.M.), Avery Huffman DIPG Foundation (to M.M.), Chadtough Defeat DIPG (to M.M.), Jacob Van de Roovaart Glioblastoma Research Fund (to M.M.), Walter V. and Idun Berry Fellowship (Y.S.K.) and Chadtough Defeat DIPG Fellowship (for Y.S.K.).

## Author contributions

S.M.G., Y.S.K., A.G., B.Y., C.M., and Y.P. designed, conducted, and analyzed experiments. R.M., A.E.I., J.R., R.D., J.H., M.Q., K.M., P.W., A.Y., and M.L, conducted and analyzed experiments. S.M.G., Y.S.K. and M.M. wrote the manuscript. J.B.Z and J.T. provided reagents and insights. M.M. conceived of the project and supervised all aspects of the work.

## Corresponding authors

Correspondence and requests for materials should be addressed to M.M.

---

**Extended Data Fig. 1.**
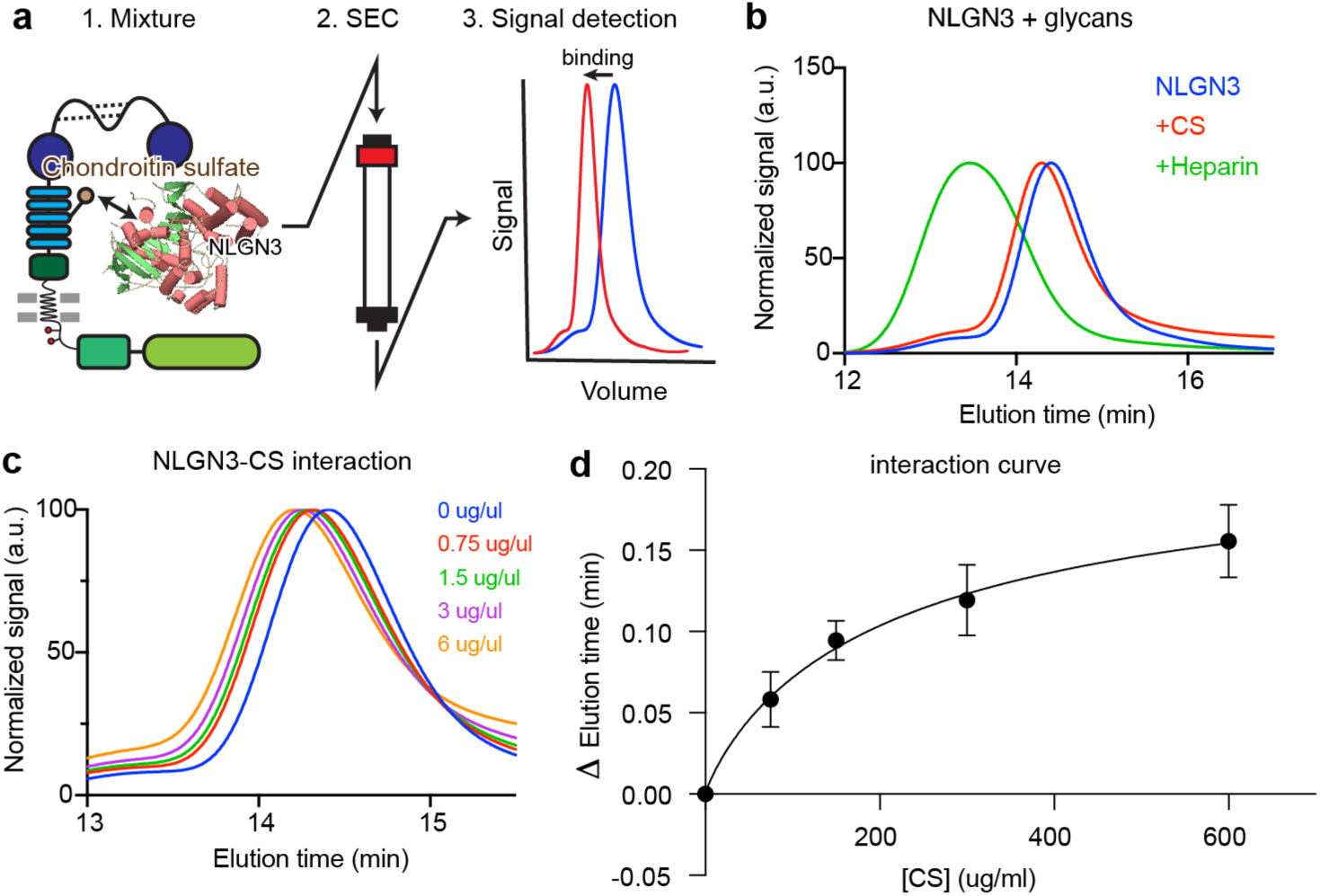
Biochemical analysis of NLGN3 interaction with Chondroitin Sulfate. **a.** Schematics of Size-Exclusion Chromatography (SEC) based coupling assay. NLGN3 and its substrates are mixed, run through the column, and the left-shift of the peak will be assessed. **b.** Binding assay of the NLGN3 with Chondroitin Sulfate (CS) and Heparin. Both Heparin and CS are binding to NLGN3, causing the peak to shift to the left. **c.** Dose-dependent interaction of NLGN3 and CS. **d.** Quantification of NLGN3-CS interaction shown in c.

**Extended Data Fig. 2.**
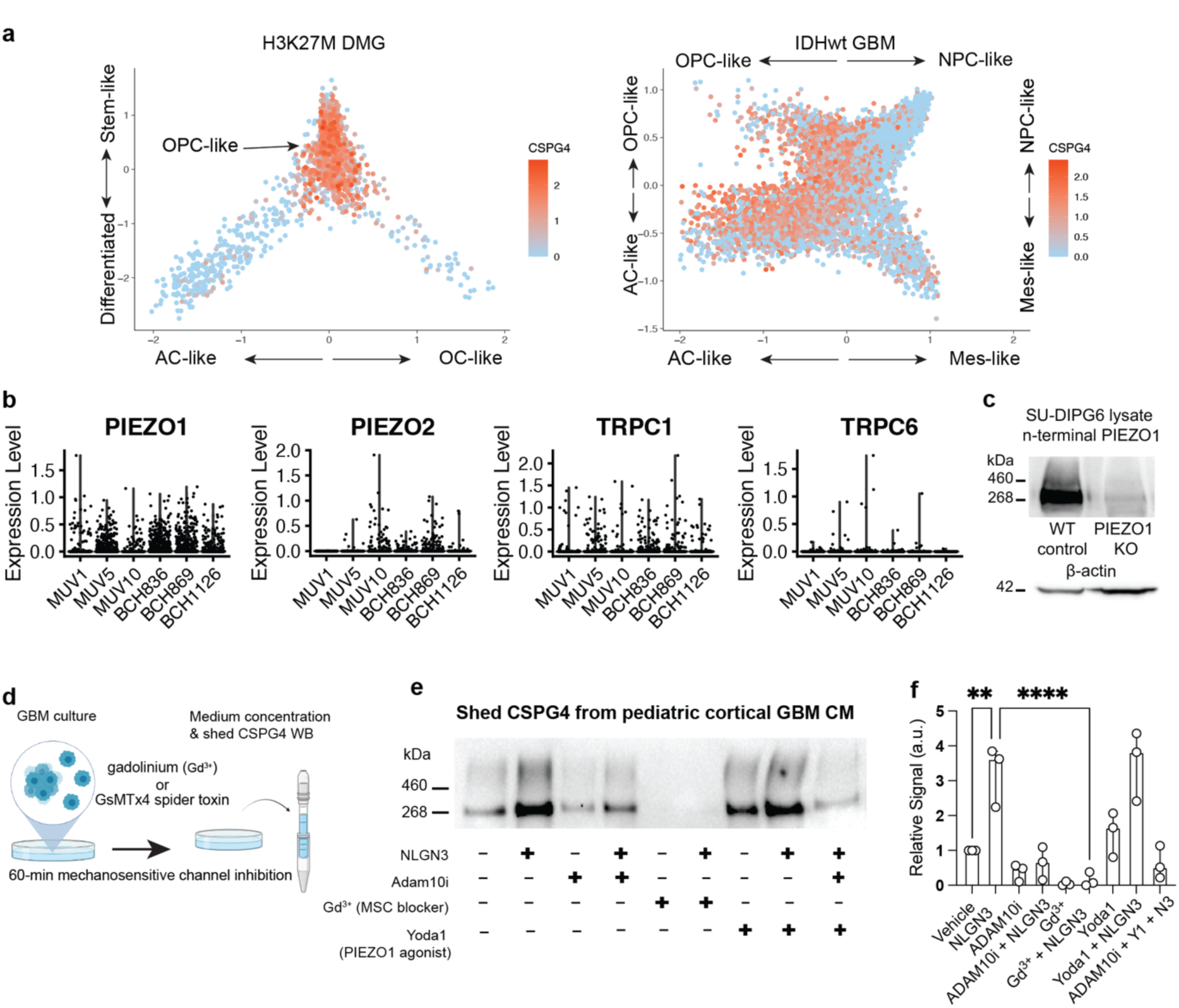
sNLGN3 induces ADAM10-dependent CSPG4 cleavage via activation of PIEZO1 channels. **a.** Scatterplots depicting tumor architecture of scRNA-seq produced from molecularly distinct patient glioma samples. From left to right are Histone H3K27M mutated diffuse midline glioma (DMG) and IDH1 wild-type glioblastoma (GBM). CSPG4 expression displayed in red across two classes of glioma. PIEZO1 and CSPG4 are both detected within the OPC-like fraction of these distinct glioma types. **b.** Violin plots of scaled gene expression level of mechanosensitive ion channels PIEZO1, PIEZO2, TRPC1 and TRPC6 in multiple patient biopsies of H3K27M mutant diffuse midline gliomas profiled by smart-seq2. All six biopsies are from Filbin et al. 2018^66^. Counts were downloaded from GEO (GSE102130) and plotted following log normalization and scaling functions in Seurat (https://satijalab.org/seurat/). **c.** Western blot of PIEZO1 protein expression from lysates of SU-DIPG6 PIEZO1 WT and KO cells. **d.** Workflow depicting treatment of pediatric cortical GBM cells with recombinant NLGN3 ectodomains in the presence or absence of inhibitors of mechanosensitive ion channels (MSC) followed by culture medium concentration and western blot to detect shedding of CSPG4. **e.** Western blot of shed CSPG4 ectodomains from pediatric cortical GBM experiments depicted in **d**. **f.** Quantification of western blots (n (independent biological replicates of western blot) = 3, mean ± s.e.m., ordinary one-way ANOVA, ** P < 0.01, **** P < 0.0001).

**Extended Data Fig. 3.**
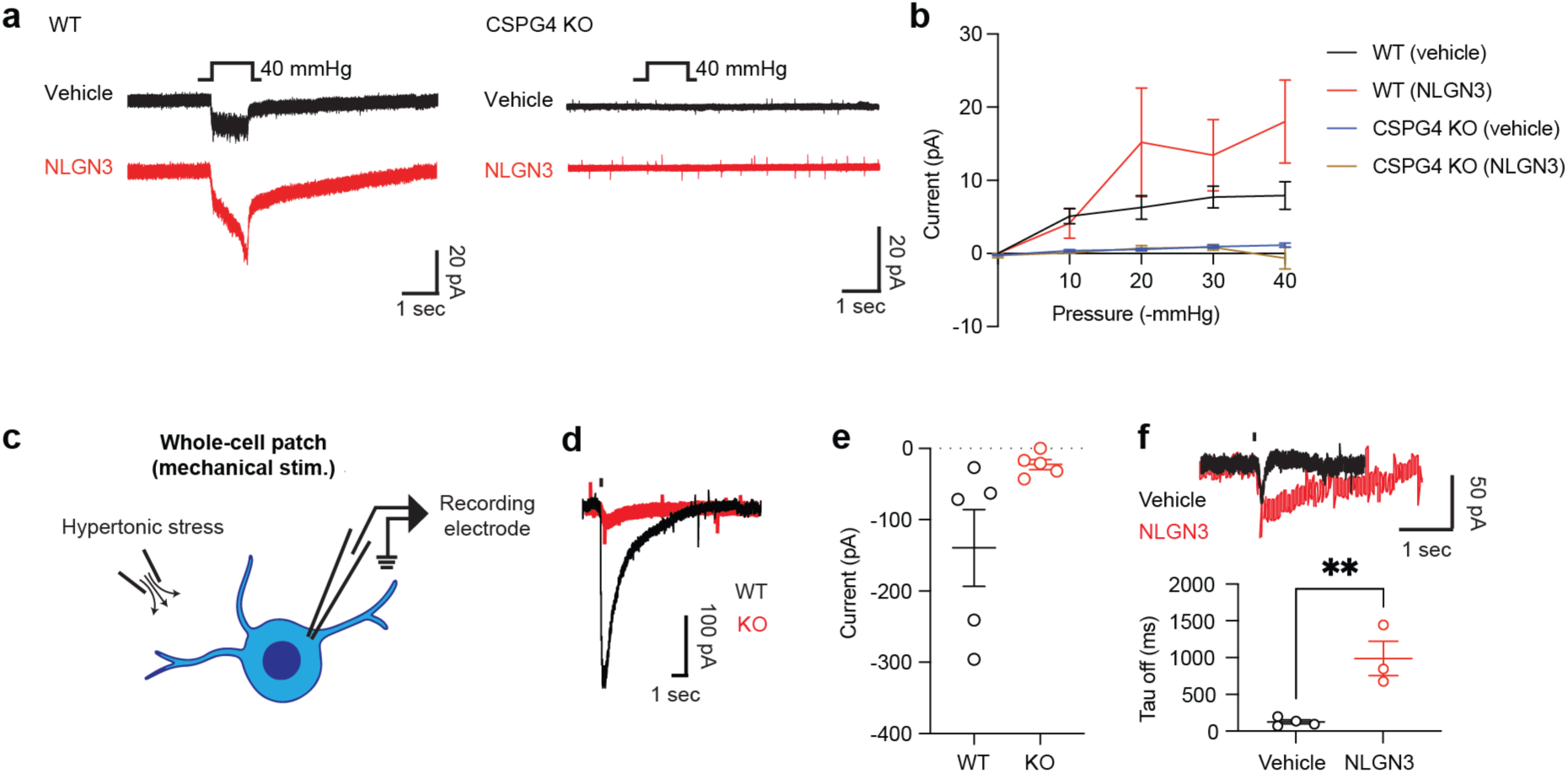
Whole-cell patch clamp characterization of PIEZO1 currents in glioma cells. **a.** Representative traces of mechanosensitive currents recorded with cell-attached recording from WT pcGBM2 glioma cells vs CSPG4 KO cells, with and without NLGN3 treatment. **b.** Summary of cell-attached experiment performed in **a** (n (number of independent cells recorded) = 5, mean ± s.e.m.,). **c.** Schematics of whole-cell patch clamp recording of mechanosensitive currents from glioma cells. **d.** Representative traces of mechanosensitive currents measured from WT and PIEZO1 KO glioma cells. **e.** Summary of the current amplitude data represented in b. (n (number of independent cells recorded) = 5, mean ± s.e.m.,) **f.** (Top) Representative currents from glioma cells with the vehicle and NLGN3 treated conditions. (Bottom) Summary of the off kinetics of the cells as shown at the top. (n (number of independent cells recorded) = 3 for the vehicle and 4 for NLGN3, mean ± s.e.m., two-tailed students t-test, ** P < 0.01)

**Extended Data Fig. 4.**
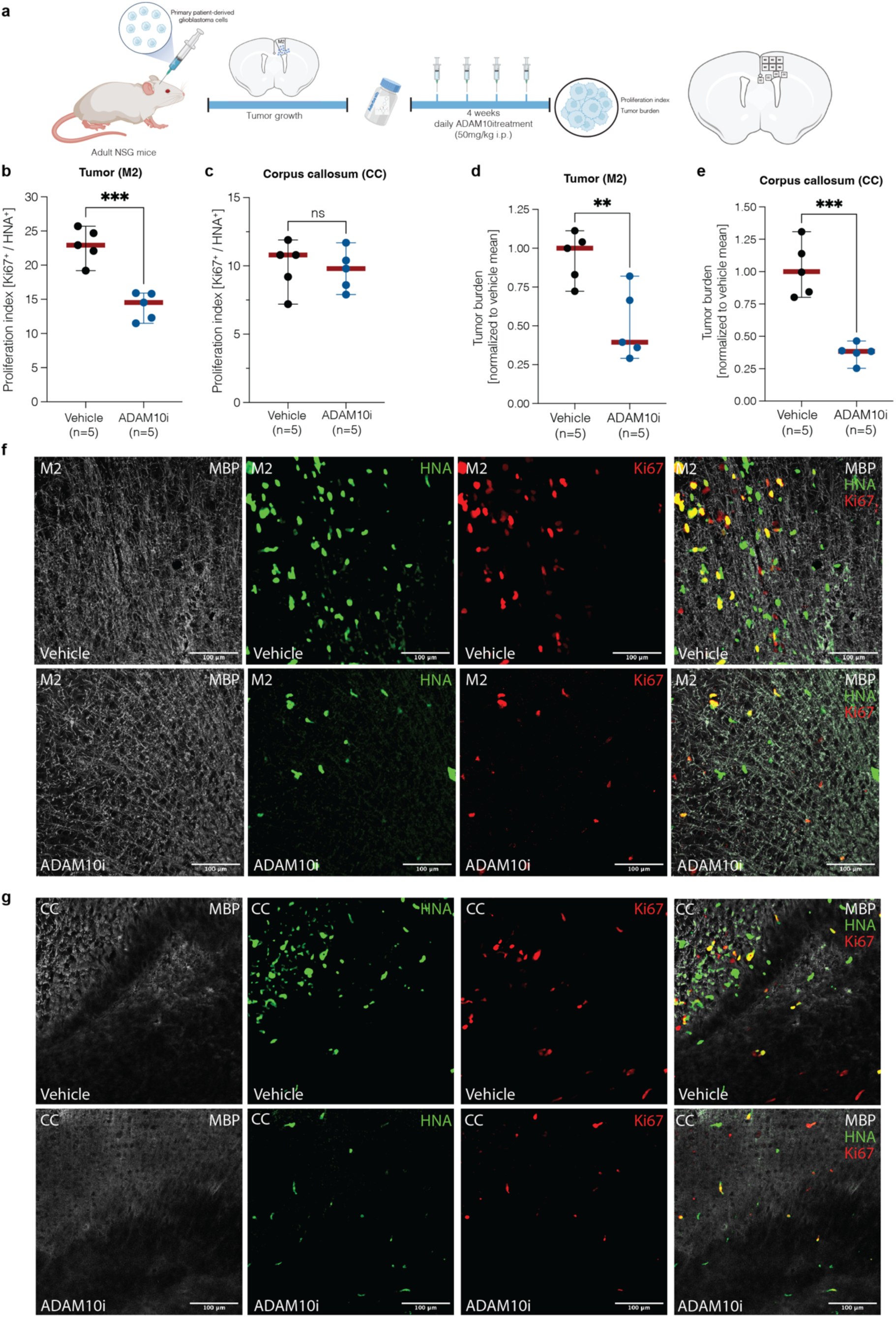
Treatment with Adam10 inhibitor decreases tumor burden in xenografted mouse model. **a.** Paradigm for *in vivo* treatment with Adam10 inhibitor aderbasib in immunodeficient mice xenografted with a patient-derived glioblastoma cell line (“*SF-232-HFC*”) and the counting model for tumor proliferation and burden. **b.** Proliferation index in vehicle- and aderbasib-treated mice (n = 5 per group) in the xenografted area of the secondary motor cortex (“M2”), quantified by confocal microscopy of Ki67^+^/HNA^+^ cells. Data are presented as mean ± range. Two-tailed student’s t-test, ***P < 0.001 **c.** Proliferation index in vehicle- and aderbasib-treated mice (n = 5 per group) of migrated glioblastoma cells into the ipsilateral corpus callosum (“CC”), quantified by confocal microscopy of Ki67^+^/HNA^+^ cells. Data are presented as mean ± range. Two-tailed student’s t-test, ns = not significant **d.** Tumor burden in vehicle- and aderbasib-treated mice (n = 5 per group) in the xenografted area of the secondary motor cortex (“M2”), quantified by confocal microscopy as the number of HNA^+^ cells. Data are presented as mean ± range. Two-tailed student’s t-test, **P < 0.01 **e.** Tumor burden in vehicle- and aderbasib-treated mice (n = 5 per group) in the xenografted area of the ipsilateral corpus callosum (“CC”), quantified by confocal microscopy as the number of HNA^+^ cells. Data are presented as mean ± range. Two-tailed student’s t-test, ***P < 0.001 **f.** Representative images of xenografted glioblastoma cells in the secondary motor cortex (“M2”) in vehicle- and aderbasib-treated mice. Gray denotes myelin sheaths labeled by MBP; green denotes HNA^+^ glioma cells; red denotes EdU. Scale bar = 100 μm. **g.** Representative images of migrated glioblastoma cells into the ipsilateral corpus callosum (“CC”) in vehicle- and aderbasib-treated mice. Gray denotes myelin sheaths labeled by MBP; green denotes HNA^+^ glioma cells; red denotes EdU. Scale bar = 100 μm.

**Extended Data Fig. 5.**
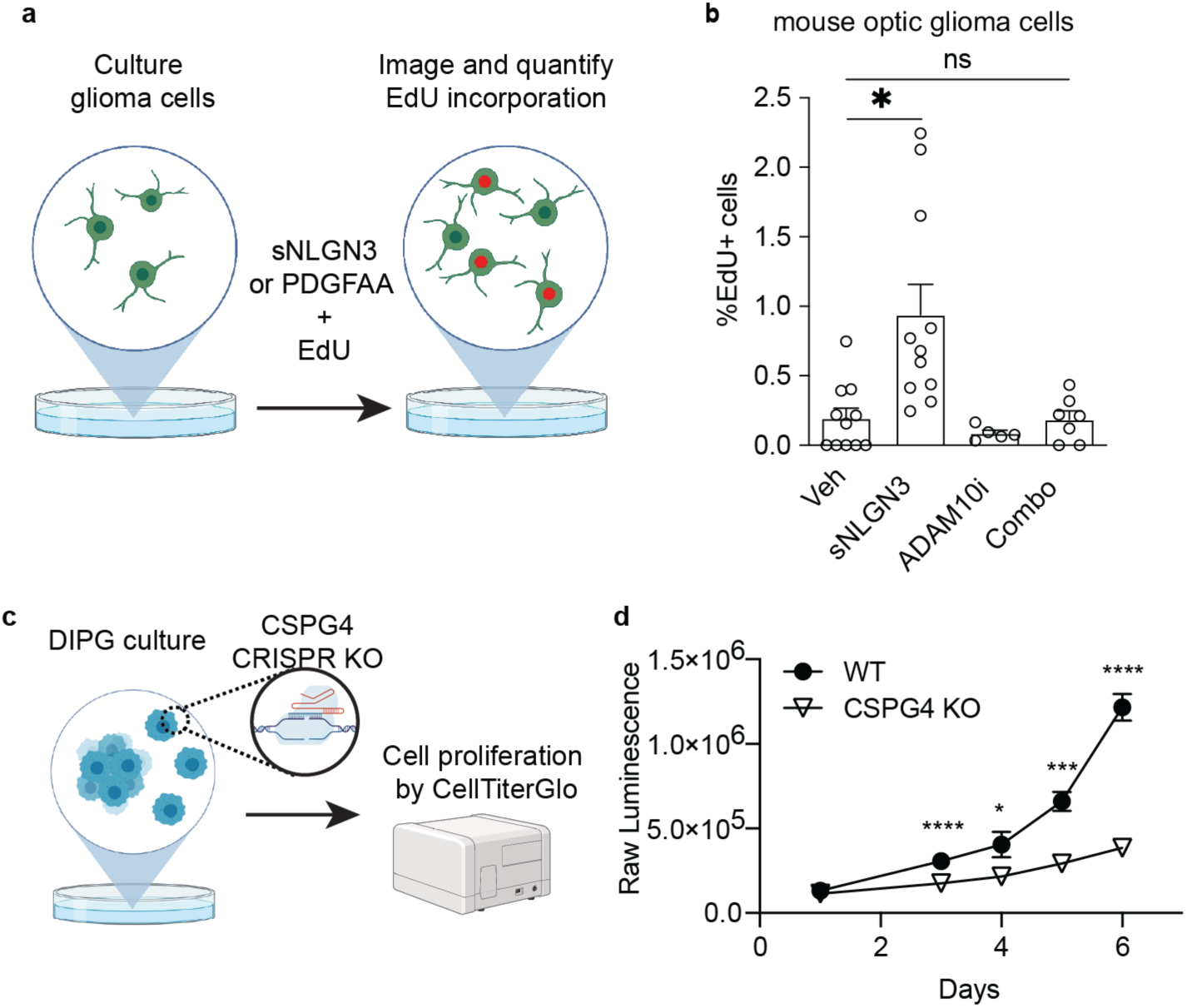
Genetic depletion of CSPG4 disrupts NLGN3-induced growth of glioma cells *in vitro*. **a.** Workflow depicting NLGN3-induced EdU incorporation assay in the presence or absence of CSPG4 shedding blocker, ADAM10i **b.** Quantification of NLGN3-induced EdU incorporation from mouse NF1-associated optic glioma cells. NLGN3 treatment increases EdU incorporation relative to vehicle controls ((n (number of independent cells) = 5-11, mean ± s.e.m., p = 0.03, one-way ANOVA with Brown-Forsythe and Welch’s test). **c.** Schematics of cell proliferation assay using CellTiterGlo assay. **d.** Quantification of CellTiterGlo proliferation assay of WT and CSPG4 KO SU-DIPGXIII- Pons cells (n (number of independent biological trials) = 3, mean ± s.e.m.,, two-tailed student’s t-test, * P < 0.05, *** P < 0.001, **** P < 0.0001). Note that from day 3 that the proliferation rate (assessed by the raw luminescence) is significantly lower in the CSPG4 KO group.

**Extended Data Fig. 6.**
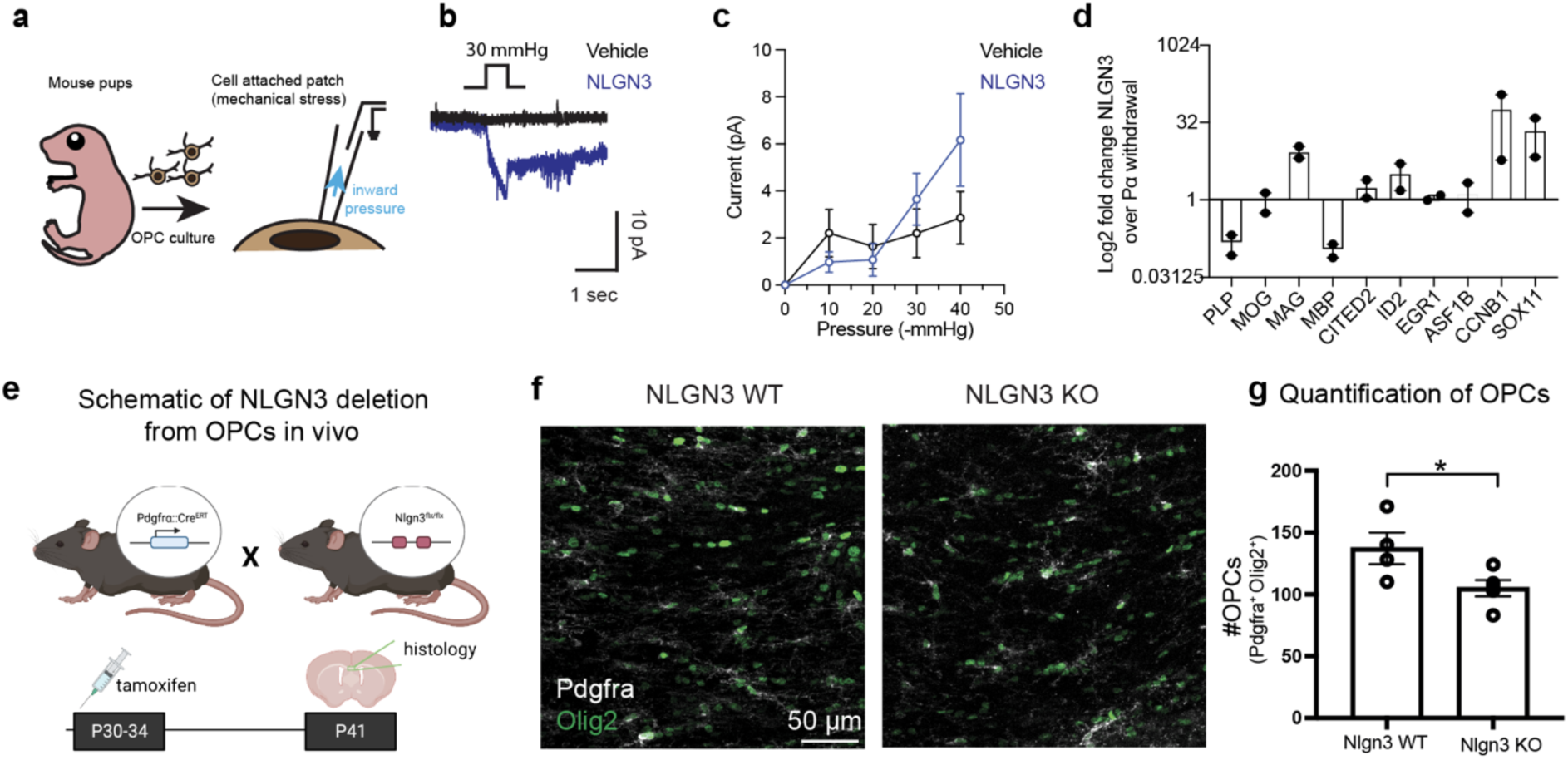
*In vivo* characterization of OPC proliferation state in NLGN3 KO mice. **a.** Schematics for *in vitro* OPC electrophysiology. **b.** Representative traces of mechanosensitive currents measured from OPCs. **c.** Summary of mechanosensitive currents shown in b. (n (number of OPCs per group) = 5 for vehicle and 6 for NLGN3, mean ± s.e.m.) **d.** qPCR results for the rat OPCs post 48 hours after PDGFα withdrawal (n (number of independent trials) = 2) with and without NLGN3 treatment. **e.** Schematics of NLGN3 deletion from OPCs in vivo. Mice expressing cell type-specific (PDGFα), inducible Cre drivers were bred to *Nlgn3*^fl/fl^ mice. Tamoxifen was administered for 5 days beginning at P21. **f.** Representative confocal images of Pdgfra positive (white) and Olig2 positive cells in WT and NLGN3 KO animals. scale bar = 50 µm **g.** Quantification of total OPC counts in WT and NLGN3 groups, as shown in **e**. (n (number of independent biological replicate of animals) = 4, mean ± s.e.m., two-tailed student’s t-test, * P < 0.05).

